# The DNA methylation landscape of the human oxytocin receptor gene (*OXTR*): Recommendations for future research

**DOI:** 10.1101/2022.03.16.484588

**Authors:** Svenja Müller, Maurizio Sicorello, Dirk Moser, Leonard Frach, Alicia Limberg, Anja M. Gumpp, Laura Ramo-Fernandez, Franziska Köhler-Dauner, Joerg M. Fegert, Christiane Waller, Robert Kumsta, Iris-Tatjana Kolassa

**Author notes:** These authors contributed equally to this work (Svenja Müller, Maurizio Sicorello, Dirk Moser). ***Correspondence and Reprint Requests:* Robert Kumsta**, University of Luxemburg, Faculty of Humanities, Education and Social Sciences, Maison des Sciences Humaines 11, Porte des Sciences L-4366 Esch-sur-Alzette, Luxemburg. E-Mail, **Iris-Tatjana Kolassa**, Ulm University, Department of Clinical & Biological Psychology, Institute of Psychology and Education, Albert-Einstein-Allee 47, 89081 Ulm, Germany.

## Abstract

The oxytocin receptor gene (*OXTR*) is of interest when investigating the effects of early adversity on DNA methylation. However, there is heterogeneity regarding the selection of the most promising CpG sites to target for analyses. The goal of this study was to determine functionally relevant clusters of CpG sites within the *OXTR* CpG island in 113 mother-infant dyads, with 58 of the mothers having experienced childhood maltreatment (CM). *OXTR* DNA methylation and gene expression was analyzed in peripheral/umbilical blood mononuclear cells. Different complexity reduction approaches were used to reduce the 188 CpG sites into clusters of co-methylated sites. Furthermore, associations between *OXTR* DNA methylation (cluster- and site-specific level) and *OXTR* gene expression and CM were investigated. Results showed that, first, CpG sections differed strongly regarding their statistical utility for research of individual differences in DNA methylation. Second, cluster analyses and Partial Least Squares (PLS) suggested two clusters consisting of intron1/exon2 and the protein-coding region of exon 3, respectively, as most strongly associated with outcome measures. Third, cross-validated PLS regression explained 7% of variance in CM, with low cross-validated variance explained for the prediction of gene expression. Fourth, very high mother-child correspondence was observed in correlation patterns within the identified clusters, but only modest correspondence outside these clusters. This study characterized the DNA methylation landscape of the *OXTR* CpG island by highlighting clusters of CpG sites that show desirable statistical properties and predictive value. We provide a Companion Web Application to guide future studies in their choice of CpG sites.

## Introduction

Both human and animal work have shown that oxytocin is involved in the development and regulation of social traits and behaviour (e.g. 1–3) as well as social cognition (e.g. 4,5). The brain oxytocin system is sensitive to environmental cues, including those acting in early developmental periods (6). It has been shown that function of oxytocin neural pathways can be influenced by early life stress (6,7), and that oxytocin receptor function across the lifespan is influenced by the degree of parental care (7). Therefore, epigenetic processes are increasingly being studied as mechanism underlying this *developmental programming* of oxytocin system regulation (8–10).

In humans, research has focused on DNA methylation in the *OXTR* promoter region, which is associated with reduced *OXTR* transcription (11). For instance, low socioeconomic status (12) as well as the degree of maternal care during childhood (13) were associated with increased *OXTR* DNA methylation in adulthood. With the focus on the effects of childhood maltreatment (CM), higher *OXTR* DNA methylation was observed in maltreated children compared with non-exposed children (14). However, some associations between childhood abuse and increased *OXTR* DNA methylation became non-significant after correction for multiple testing (15,16) and null-findings were also reported (17). A recent longitudinal investigation of mother-infant dyads showed that low maternal engagement was associated with increased variance of infant saliva-derived *OXTR* DNA methylation from five to 18 months, indicating reductions and increases over time (18). This study provided first evidence that *OXTR* DNA methylation might be dynamic in children but relatively stable in motherhood (18). With regard to further investigations in mother-infant dyads, one study reported significantly higher overall *OXTR* DNA methylation in mothers with persistent perinatal depression (PD) (19), while another study observed significant associations between CM experience and altered site-specific *OXTR* DNA methylation in mothers (20). However, no differential *OXTR* DNA methylation was detected in children of mothers with PD or CM experience (19,20). Finally, a recent meta-analysis comprising 15 independent samples concluded that early adversity is related to higher *OXTR* DNA methylation levels (8). However, the effect was very small (r=.02) and only significant in the set of studies using nonclinical samples or in those that assessed DNA methylation in blood samples (8). Nevertheless, as the authors note, there is large heterogeneity between these studies regarding methodology and genomic regions that were analyzed, which impedes meta-analytic inference (8).

Transcription of the *OXTR* is controlled by a promoter whose CpG island constitutes 2519 basepairs including 188 CpG sites (11,21), representing a large search space for possibly functionally relevant CpG sites. The majority of studies targeted single or only a few (about 20) individual CpG sites either in the so-called MT2 area (e.g. 14,18) containing 27 CpG sites with putative functional relevance (11) or in the protein-coding region of exon 3 (e.g. 13,19) (see 17,20 for exceptions; see Supplementary Table 1 for a detailed overview of previously investigated CpGs sites). This heterogeneity in selecting CpG sites within the *OXTR* CpG island reflects a lack of consensus concerning the best sites to target for DNA methylation analysis (8).

The current study thus aimed at determining functionally relevant clusters of CpG sites within the *OXTR* CpG island in mother-infant dyads. The entire CpG island comprising 188 CpG sites was analyzed with targeted deep bisulfite sequencing (22). Different complexity reduction approaches were used to reduce the amount of 188 CpG sites into functionally relevant clusters of co-methylated CpG sites. Furthermore, the association between *OXTR* DNA methylation (on level of cluster and single CpG sites) and *OXTR* gene expression levels as well as CM experiences were investigated.

## Material and Methods

### Participants and study procedure

Women who gave birth at the maternity ward of the Ulm University Hospital between October 2013 and December 2015 were invited to participate in the project *My Childhood* – *Your Childhood*. Exclusion criteria for study participation were age under 18 years, insufficient knowledge of the German language, and severe health problems of mother or child during pregnancy or labour. Out of 5426 mothers invited, 533 mothers gave written informed consent and sociodemographic characteristics were assessed. Immediately after parturition, 45 ml umbilical cord blood was collected into CPDA-buffered tubes (Sarstedt S-Monovette, Nürmbrecht, Germany). Within in the following week, peripheral blood was drawn from the mothers.

Maternal childhood maltreatment (CM) exposure was assessed with the German short version of the *Childhood Trauma Questionnaire* (CTQ; 23) including five subscales (i.e. emotional, physical and sexual abuse, emotional and physical neglect). Using a specific cut-off (24), mothers with mild to severe CM experiences in at least one CTQ subscale were categorized as CM+. Mothers without a history of CM were categorized as CM-. The epigenetic analyses were conducted in a subset of study participants including 117 mothers (*n*=59 CM-, and *n*=58 CM+) and 113 infants (for subset selection and more details of the cohort see (20)). The study followed the principles of the Declaration of Helsinki and was approved by the local ethics committee of Ulm University.

### DNA methylation analysis

DNA was isolated from peripheral blood mononuclear cells (PBMCs) from the mothers and umbilical blood mononuclear cells (UBMCs) from the newborns as described elsewhere (20). Sodium bisulfite conversion of 500 ng of genomic DNA from each subject was performed using the EZ DNA Methylation-Gold^™^ Kit (Zymo, USA). DNA methylation levels at specific genomic sites were investigated by targeted deep bisulfite sequencing following the protocol by Moser et al. (22). We focused on regulatory regions of the *OXTR* gene: A CpG island comprising 188 CpG sites (11,21) and 21 CpG sites at a potential enhancer element in intron 3 (15) (for detailed description of chromosomal positions see Supplementary Table 1). The following pair-end sequencing was performed on a MiSeq system (Illumina; San Diego – USA) using the Illumina MiSeq reagent Kit v2 (500 cycles-2×250 paired end) in collaboration with the BioChip Labor of the Center of Medical Biotechnology (ZMB, University Essen/ Duisburg). See Supplementary Material for gene-specific primer sequences (Supplementary Table 2) as well as quality controls in terms of mean sensitivity (Supplementary Table 3) and standard curves to demonstrate linearity (Supplementary Figure 1) for the different assays.

For sequence analysis, FASTQ files from all sequencing reads were generated to enable software analysis using amplikyzer2 (25,26). Quality control led to the exclusion of seven CpG sites in genomic regions of two overlapping amplicons that did not show perfectly matched DNA methylation values (CpG 73-77; 87,88; see Supplementary Table 1 for chromosomal positions). The excluded CpG sites showed low DNA methylation values (*M*=2.43%) and low variability (*SD*=1.44%). CpG sites influenced by *OXTR* single nucleotide polymorphisms (introduction, disruption or shift in a CpG site) were excluded from analysis (CpG 8; 15a; 59; 72a; 83; 92; 128; E13). In addition, one CpG located in primer binding region (enhancer region) was excluded from statistical analysis. Thus, the final dataset for statistical analyses consisted of 183 CpG sites in the *OXTR* promoter and 19 CpG sites in the enhancer region. Furthermore, samples with missing values in at least one or more CpG sites were excluded, which resulted in the exclusion of seven mothers and thirteen infants. Thus, the final dataset comprised 110 mothers and 100 infants.

### Gene expression analysis

Gene expression data was available from a previous study by Ramo-Fernandez et al. (20). In brief, total RNA from a subset of participants (67 mothers and 34 infants) was extracted from PBMC/UBMP using the Qiagen RNeasy Kit (QIAGEN, Hilden, Germany). Subsequently, cDNA synthesis was conducted using the cDNA reverse transcription kit (Thermo Fisher Scientific, Germany) followed by real-time quantitative PCR analyses of the *OXTR* on a QuantStudio 6 qPCR platform (Life Technologies, USA) using the appropriate Taqman gene expression array (Hs00168573_m1; Thermo Fisher Scientific). According to the program NormFinder (27), succinate dehydrogenase complex, subunit A, flavoprotein variant (*SDHA*) and importin 8 (*IPO8*) were selected as gene expression references. Relative mRNA expression levels of *OXTR* were defined with the 2-ΔCt equation, with ΔCt = (mean Ct of the target) – (geometric mean of the Ct of the reference genes SDHA and IPO8). The resulting fold-change values – an estimate of relative mRNA expression levels – were used for statistical analyses.

### Statistical analysis

#### Cluster analysis

Two complementary approaches were used to identify data-driven clusters of similarly methylated CpGs. First, hierarchical agglomerative clustering was performed on a matrix of Euclidean distances using Ward’s method. The number of clusters was identified based on the elbow method for within-cluster sums of squares and the silhouette method using the R package factoextra (v1.0.7). Second, an algorithm called Density-Based Spatial Clustering of Applications with Noise (DBSCAN) was used with the R package DBSCAN (v1.1.8; 28). While the hierarchical agglomerative clustering approach clusters all data points until all are in the same cluster, DBSCAN only clusters data points based on two prespecified hyperparameters (minPts, the minimum cluster size, and ε, the radius around a given data point that is searched for other points). This allows distinguishing between clusters of CpG sites and single CpG sites that do not belong to a coherent cluster, which are summarized into a “noise” category. The hyperparameter ε was chosen by selecting different values for minPts (3, 5, 8, 12) and identifying the knee in each plot of k-nearest-neighbour distances between all points. The y-axis value at the knee is the chosen hyperparameter ε. Both clustering approaches were performed on z-standardized data.

#### Partial Least Squares prediction of mRNA expression and trauma

In the subsample of mothers, the variance at each CpG site was compared to the assay sensitivity on this site using a chi^2^ test and 109 degrees of freedom. CpG sites were excluded from machine learning analysis if their variance was not significantly larger than the assay sensitivity. We used Partial Least Squares (PLS) to identify latent components in the CpG methylation data which are maximally predictive of mRNA expression, CTQ scores, and trauma versus control group membership. The PLS algorithm has one hyperparameter, which is the number of latent components that should be extracted. This hyperparameter was tuned in a 5-fold inner cross-validation loop and the performance of the resulting model assessed using a 5-fold outer cross-validation loop. This procedure was repeated two times to counteract the effect of random fold slicing, leading two a 2×5×5 cross-validation procedure. For mRNA expression and CTQ, PLS regression was performed with rootmean-squared error (RMSE) as the performance metric for parameter tuning, while R2 is reported as it is easier to interpret for most readers. For trauma group membership, PLS discriminant analysis was performed with accuracy (percentage of correct classifications) as the performance metric. Significance of the models was assessed by randomly permuting the respective outcome 1000 times and calculating the proportion of performance metrics below (RMSE) and above (accuracy) the empirical (non-permuted) value. To inspect how single CpGs load on the identified latent components, we refitted PLS models with the optimal number of components without cross-validation. PLS was performed using the R packages caret (v6.0-90; 29) and pls (v2.8-0; 30).

#### Multivariate mediation

We tested whether CpG sites can be clustered according to their propensity to mediate the effect of traumatic experiences on mRNA expression in mothers. To this end, we employed the principal directions of mediation (PDM) approach in Matlab (2020a), which has been recently developed and tested in neuroimaging applications to identify mediating brain networks (31,32). PDMs are orthogonal latent components on which the CpG-wise DNA methylation values can be projected. These components are chosen to maximize the mediation effect of trauma on mRNA expression via CpG methylation. Similar to principal component analysis, the first component will mediate the largest portion of the indirect effect, followed by the second component and so on. This method can be understood as an extension of PLS to mediation models.

First, the dimensionality of DNA methylation data was reduced to 20 components using single value decomposition in correspondence with the method developers, as using all single CpG sites led to unstable results. By default, 5 PDMs are built. Plotting the indirect effect mediated via each component can be used to identify how many PDMs should be extracted, analogous to the scree plot in principal component analysis.

#### Analysis of differentially methylated positions and regions

In addition to cluster- and component-based analyses, we also examined 202 CpGs independently to identify differentially methylated positions (DMPs) and differentially methylated regions (DMRs) in association with group status, CTQ and *OXTR* mRNA expression in the subsample of mothers. To test for DMPs, we used the R package limma (v3.46.0; 33) and further conducted separate t-tests or linear models for each CpG as sensitivity analyses. For the analysis of DMRs, we used the R package DMRcate (v2.4.1; 34) to identify regions of CpG sites within a range of 500 base pairs (bp) in association with the respective measure of interest, and the R packages edgeR (v3.32.1; 35) and bsseq (v1.26.0; 36) for preparation of the DNA methylation data. Further exemplary sensitivity analyses included adjustment for maternal age and varying the bp range in the DMR analysis.

#### Script Accessibility

All analysis scripts can be accessed on: https://github.com/MaurizioSicorello/OXTRmeth_analyses

## Results

### Qualitative description of *OXTR* methylation

Sections of the *OXTR* gene differed strongly in their statistical adequacy for research on individual differences in DNA methylation (Figure 1). The section of exon 1, which lies upstream of the MT2 region, had extremely low DNA methylation levels and dispersion, while also being characterized by a considerable number of outliers (Mdn = 0.00, IQR = 0.01, mean skew = 4.61). Both median and IQR were well below the median sensitivity of the assay across all CpGs (Mdn = 0.02, IQR = 0.1, range = 0.0002-0.07). In contrast, CpGs in the MT2 region exhibited desirable statistical properties for clinical studies, including higher median levels, higher dispersions, and symmetrical distributions (Mdn = 0.10, IQR = 0.15, mean skew = 0.58). These desirable properties extended to an early section of intron 1 outside the MT2 region. Then, within intron 1, median levels and dispersions dropped again to very low levels, a pattern extending into downstream sites including exon 2, intron 1, and roughly 10% of exon 3, albeit with low levels of outliers compared to exon 1. A next section within exon 3 was characterized by maintained low median levels, but a marked number of extreme outliers. A last large section of exon 3 had desirable statistical properties, including higher median levels and dispersions. The same qualitative pattern was found in children (Supplementary Figure 2). For sections of the potential *OXTR* enhancer element see Supplementary Figure 3A (mothers) and Supplementary Figure 3B (children).

**Figure 1:**
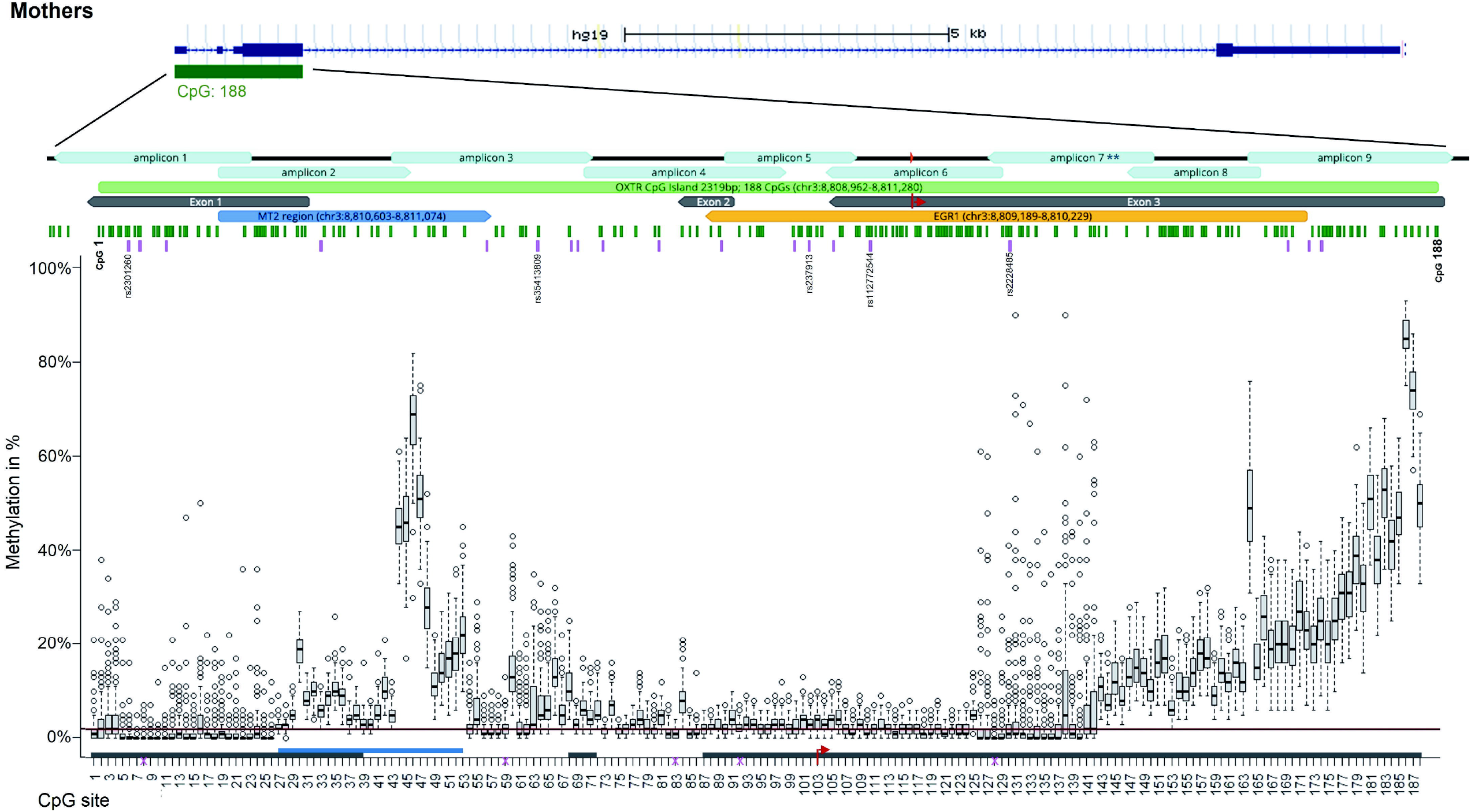
Chromosomal position of OXTR promoter region with corresponding DNA methylation levels (mothers). Chromosomal position of all fragments of the OXTR CpG island for mothers are illustrated using graphical outputs generated by the UCSC Genome Browser (https://genome.ucsc.edu) and Geneious Prime 2020 software (https://www.geneious.com). The gene is presented in 5’-> 3’ orientation from left to right. Amplicons are highlighted with respect to their genomic orientation. All CpG sites are illustrated with green bars, CpG island in light green (with CpG1 and CpG 188 labeled), MT2 area in blue, and exons in grey colour (transcription start site is marked by the red arrow). A region including several transcription factor binding sites for EGRI (OREG1492306 according to the UCSC Genome Browser; also known as NGFIA) is indicated in yellow. Common SNPs with a minor allele frequency >5% according to the UCSC Genome Browser are illustrated with pink bars. Genomic regions of OXTR SNPs that introduce, disrupt or shift methylation sites are highlighted (these CpG sites were excluded from statistical analyses). Boxplots showing DNA methylation across the investigated CpG sites. The box covers the DNA methylation data of each CpG site between the 25th to 75th quantile, the whiskers show the range of values falling within 1.5-fold the interquartile range. The horizontal line (red) represents the DNA methylation detection limit for targeted deep bisulfite sequencing (*OXTR* promoter: 1,96%). Note: ** The values in the range of amplicon 7 are not different due to epigenetic heterogeneity but rather due to technical variability.

For CpG-wise summary statistics, e.g. univariate statistical features and associations with childhood trauma and *OXTR* mRNA expression, including flexible thresholds and Bayes factors, see Supplementary Table 1 or the web application.

### Correlation of methylation between CpG sites

Visually, there was a clear cluster of highly correlated CpG sites within exon 3 (Figure 2A/B; CpGs 143-193 in both mothers and children). The average correlation within this block was *r*=.76 for mothers and *r*=.81 for children, demonstrating substantial redundancy. Correlations were also very high in a subsection of intron 1 (CpG 53-65), with average *r*=.79 for mothers and *r*=.80 for children, albeit the limit of the cluster was visually less clear and could also comprise sections of exon 2. Overall, there was high correspondence between correlation patterns in mothers and children (*r*=.75), but visual inspection revealed this large effect was mainly due to extremely high mother-child correspondence in the exon 3 (*r*=.93) and intron 1 (*r*=.87) clusters described above, while the CpG correlation patterns outside these clusters exhibited only modest mother-child correspondence (*r*=.32). This suggests a high generalizability of correlation patterns within these two clusters for different developmental stages (adult *vs*. child) and different tissues, while correlation patterns outside these regions exhibit low generalizability.

**Figure 2.**
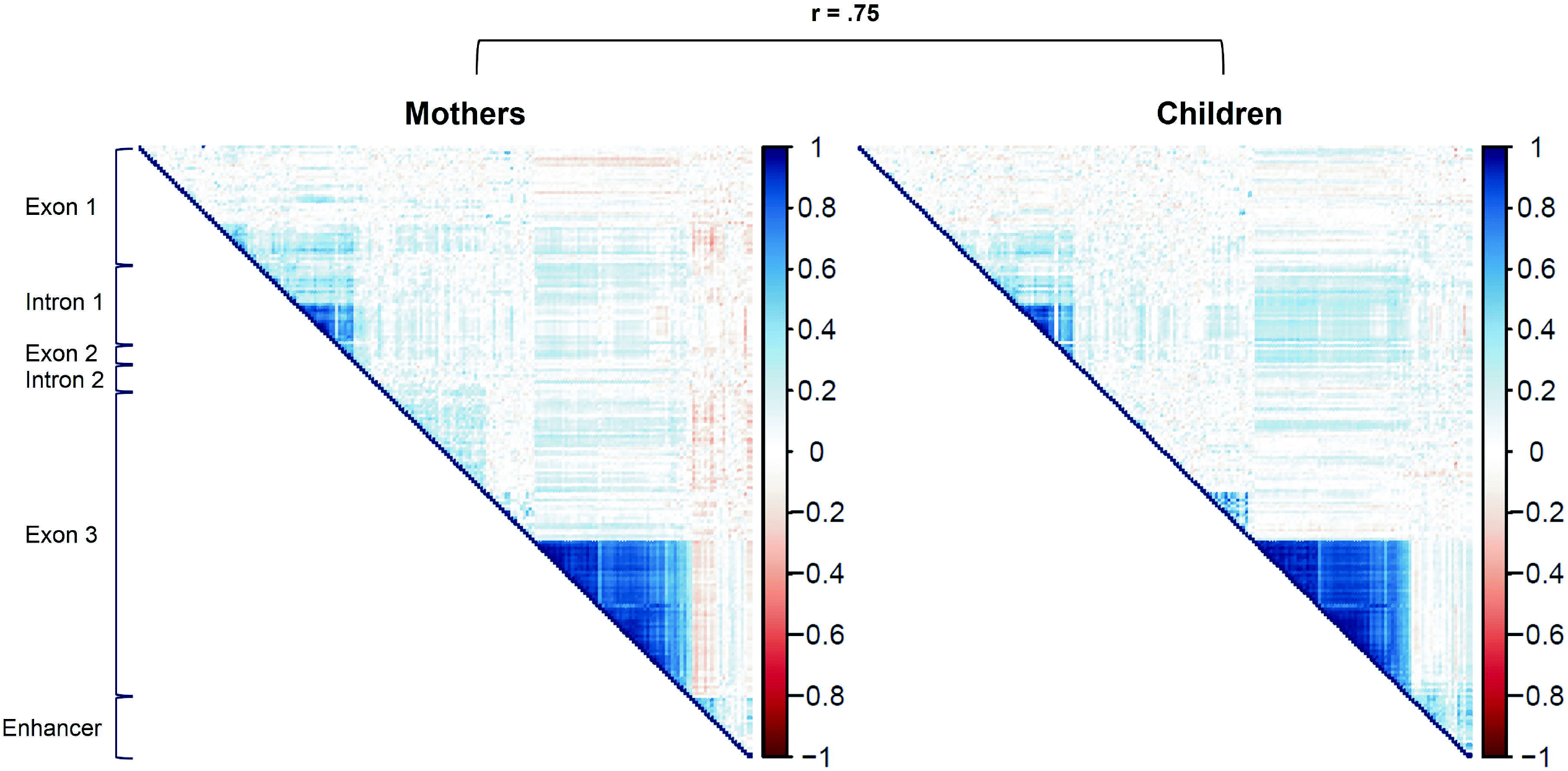
Correlation between *OXTR* DNA methylation at different CpG sites for mothers and children. All CpG sites of the *OXTR* CpG island (comprising three exons and two introns) and a potential enhancer element were included. In addition, the correspondence between correlation patterns in mothers and children is shown.

### Cluster analysis

The results of the agglomerative hierarchical clustering analysis are shown in Figure 3. The silhouette and elbow method suggested the existence of three clusters. One cluster consisted entirely of the protein-coding section in exon 3, for which we already observed high within-cluster correlations (blue dendrogram cluster, CpGs 142-192; see section above). A second cluster consisted of adjacent CpG sites of intron 1 (74%) and exon 2 (26%; yellow dendrogram cluster). The complementary DBSCAN clustering approach also suggested a three-cluster solution that assigned 92% of data points to the same clusters as the hierarchical clustering procedure and remained stable when different hyperparameters were used (Supplementary Figure 4). Importantly, the DBSCAN procedure suggests the third cluster is an outlier category for data points which cannot be sensibly clustered. Hence, taken together, visual inspection of correlations, hierarchical clustering, and DBSCAN suggest two clusters mainly consisting of intron1/exon2 segments and the protein-coding segment of exon 3, respectively. Comparing these clustering solutions to solutions based on the child sample showed an overlap in CpG-to-cluster assignment of 98% for the hierarchical clustering approach and 100% for DBSCAN (Supplementary Figures 5-6).

**Figure 3.**
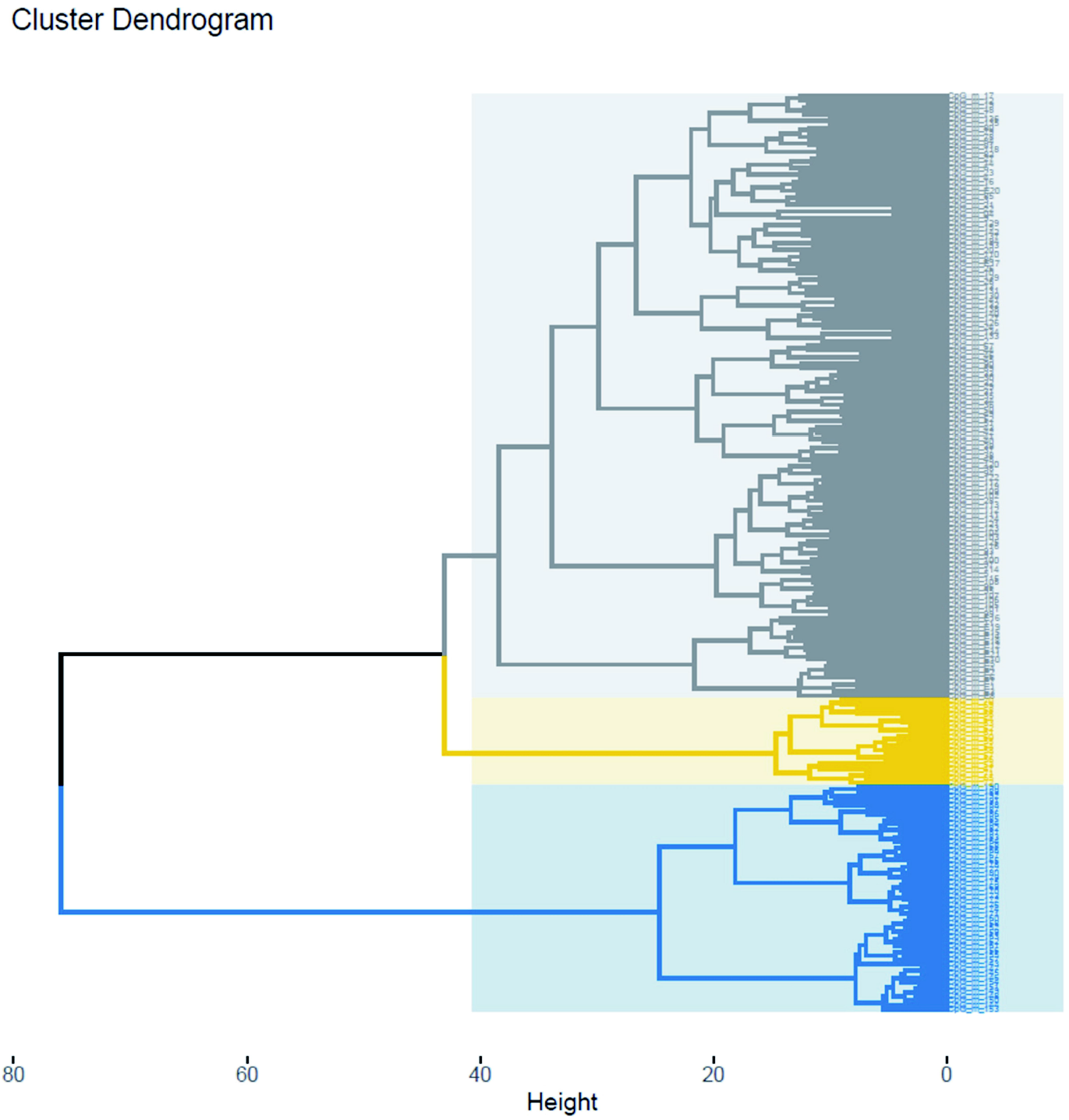
Hierarchical agglomerative clustering solution for *OXTR* DNA methylation in mothers. All CpG sites of the *OXTR* CpG island (comprising three exons and two introns) and a potential enhancer element were included.

### Clustering based on associations with childhood maltreatment and mRNA expression

Thirty-six CpG sites were excluded due to insufficient variance, leaving DNA methylation of 166 CpG sites as predictors. The cross-validated PLS regression explained 8.74% of variance in *OXTR* gene expression, which was statistically not significant (*p*=.092). Variance explained for continuous CTQ scores was 7.00% and statistically significant (*p*=.040), albeit the model with trauma as a binary outcome did not yield a statistically significant cross-validated classification accuracy (55.60%, *p*=.122). For all three models, the cross-validated performance was optimal with only one latent component (Supplementary Figure 7-9). The loadings of DNA methylation at single CpG sites on the latent components predicting gene expression and CTQ show that the clusters identified above (intron1/exon2 and protein-coding section of exon3) contribute most strongly to this latent dimension together with several CpG sites within MT2 (Figure 4). For the binary trauma outcome, the same result was found for the exon 3 section, while the other two sections did not contribute (Supplementary Figure 10). Notably, we repeated the PLS regression procedure with outcome variables created from a random normal distribution. The results indicate that high intercorrelations within the identified clusters might contribute to their high loadings, even in the absence of a meaningful external criterion (Supplementary Figure 11).

**Figure 4.**
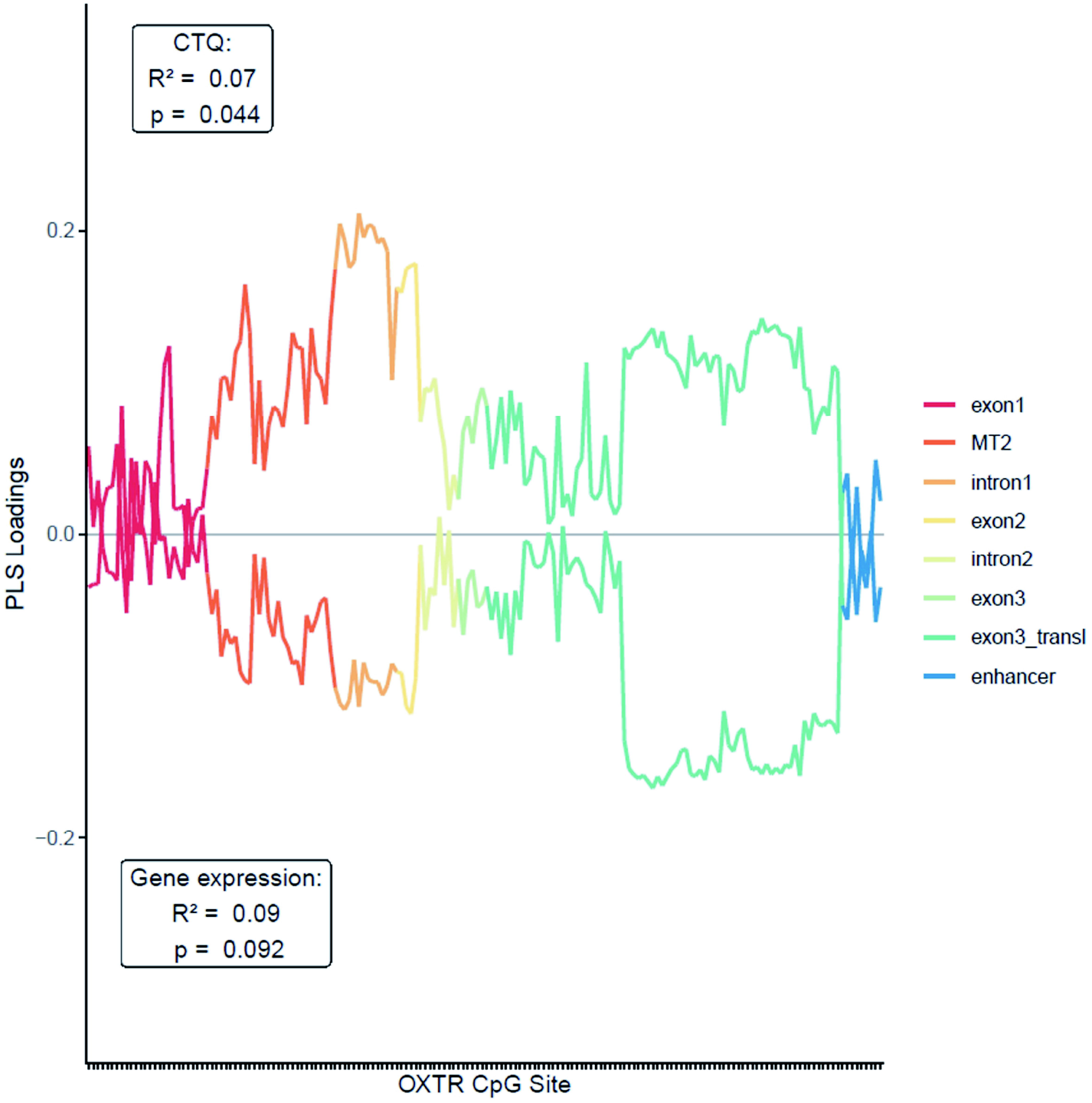
Partial Least Squares loadings of *OXTR* DNA methylation at single CpG sites for the prediction of childhood maltreatment and *OXTR* mRNA expression. Analyses included 166 CpG sites of the OXTR CpG island as predictors, after excluding thirty-six CpG sites due to insufficient variance. The upper line corresponds to a model predicting scores on the childhood trauma questionnaire (CTQ). The lower line corresponds to the model for mRNA expression levels.

### Multivariate Mediation

For the mediation of CTQ on gene expression via DNA methylation patterns, the scree plot for the mediation path suggested up to four mediating components. The mediation path via the first component had a small-to-moderate effect size, but the confidence intervals were extremely large, indicating that much larger samples are needed to provide sufficiently precise estimates with this method (mediation path = -.25, 95% CI = [-.75, .30]). Similar results were found for binary trauma as the predictor (mediation path = -.27, 95% CI = [-.68, .46]).

### Analysis of differentially methylated positions and regions

The analysis of single CpG sites in association with CM using the group status of 110 mothers revealed three significant DMPs after correcting for multiple testing (FDR<.05; Supplementary Table 4), which are located in the protein-coding section of exon 3 (CpGs 172, 173 and 177). Results somewhat differed using the CTQ scores as measure of CM (Supplementary Table 5), showing one significant DMP located in exon 1 (CpG 18) and one in the protein-coding section of exon 3 (CpG 135). For the association between DNA methylation and mRNA expression of *OXTR* (*N*=69), we did not find any significant DMPs (Supplementary Table 6). Results were comparable when adjusting for maternal age or when using *t-tests* (CM group status) or linear models (CTQ/mRNA expression) instead of the limma package (Supplementary Tables 7-11).

Analyzing regions of adjacent CpG sites (Supplementary Table 12) revealed a significant DMR in association with CM using the group status, including nine CpGs in the protein-coding section of exon 3 (CpGs 168-176). The number of CpGs within the DMR however, was sensitive to variation of ranges between CpG sites other than the default parameter of DMRcate (500bp), including between four (250bp) and 16 CpGs (750bp), and also varied when adjusting for maternal age (13 CpGs). Using CTQ as measure of CM, we found a significant DMR covering four CpGs in intron 1 (CpGs 61-64). We did not find any significant DMRs in association with *OXTR* mRNA expression.

Associations (-log10 p-values) of DNA methylation at each CpG site with CTQ and with mRNA expression are depicted in Figure 5, and associations of DNA methylation of all CpGs with group status and mRNA expression are shown in Supplementary Figure 12.

**Figure 5.**
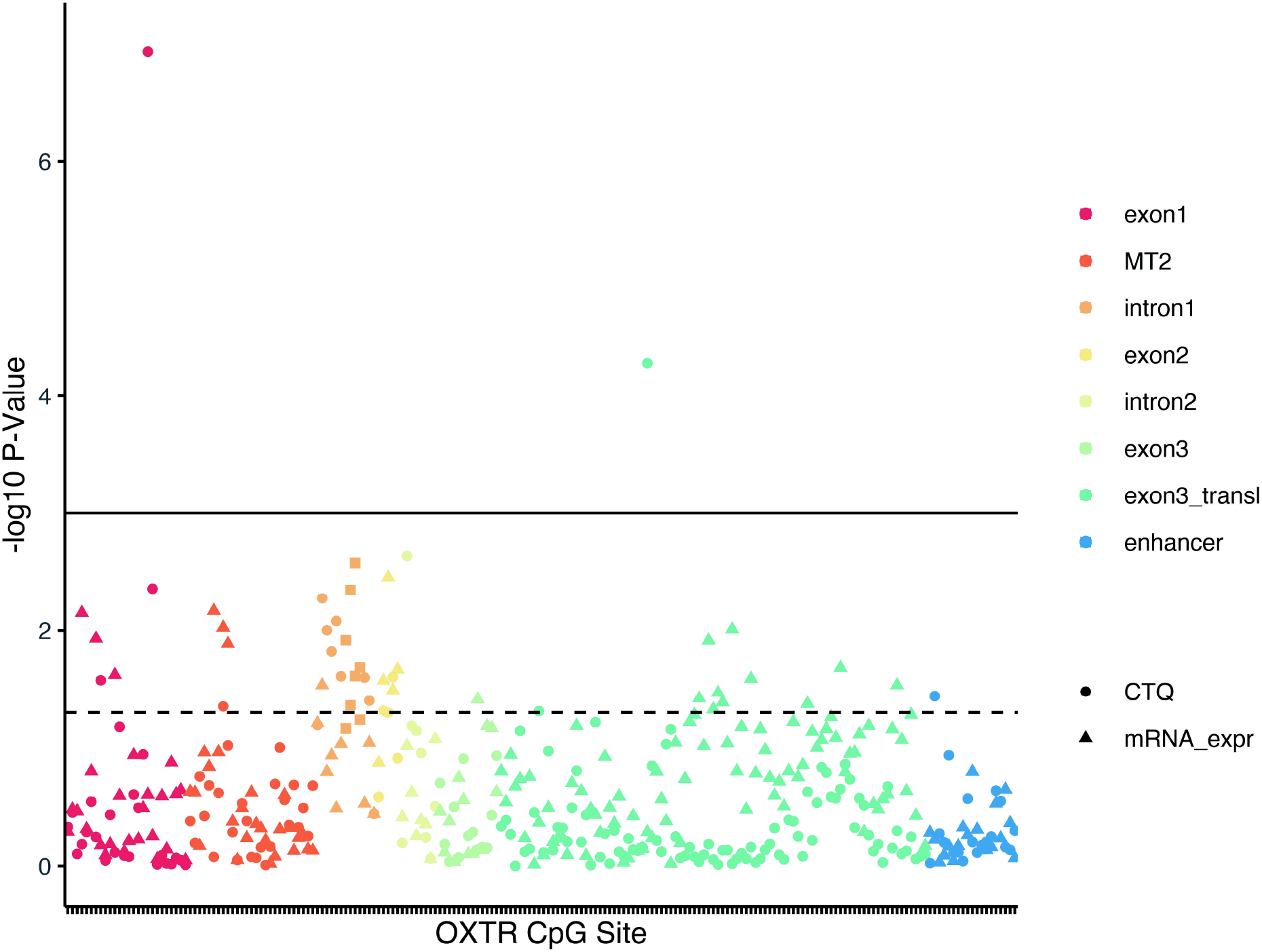
Associations of childhood maltreatment (CTQ) and *OXTR* mRNA expression with *OXTR* DNA methylation at single CpG sites. Triangles depict the-log10 p-value of the association between DNA methylation and mRNA expression, circles depict the association with childhood maltreatment (CM) and filled squares indicate the DMR in association with CM. The dotted line indicates a nominal p-value of .05, whereas the solid line indicates an FDR of .05.

Curves depicting statistical power as a function of effect size, sample size, and significance threshold can be found in Supplementary Figure 13-14.

### Associations with average *OXTR* DNA methylation across CpG sites

To explore the utility of a simple baseline model, we averaged the *OXTR* DNA methylation for each participant in the mother subsample across all CpG sites. This approach has been previously applied in other studies (20), although in the most cases averaging was performed over a substantially smaller number of CpG sites (e.g. 37). These average scores were significantly correlated with trauma group assignment (*r*(108)=.25, *p*=.009), gene expression (*r*(67)=-.28, *p*=.018), but not CTQ score (*r*(108)=.15, *p*=.111). The significant associations survive Bonferroni-Holm correction for three comparisons. Still, these in-sample statistics cannot be compared to the out-of-sample accuracies of the predictive machine learning models above, which are expected to be lower. Predicting the three outcomes from DNA methylation in separate univariate linear models using 5-fold cross-validation with 100 repeats resulted in negative R2 values in all three cases, granting no evidence for generalizable predictive utility. As the number of amplicons is the limiting financial factor for future studies, we report the correlations between the three external outcomes and average DNA methylation within each amplicon in the supplements (Supplementary Figure 15).

### Companion Web Application

We designed a shiny web application to guide researchers in the decision which CpG sites to investigate in their research: https://msicorello.shinyapps.io/oxtr-shinyapp/. It allows selecting CpG sites based on the data presented here and several user-defined filters, e.g. their chromosomal location on the gene, univariate statistical features, and associations with the three external outcomes, including flexible thresholds and Bayes factors. A table of selected CpG sites can be downloaded from the app. Besides CpG-wise summary statistics, this table also includes additional information, for example, whether CpG sites are covered on Illumina Infinium Arrays (450k/EPIC) or in previous peer-reviewed publications on early adversity.

## Discussion

There is growing interest in the investigation of *OXTR* DNA methylation in humans, accompanied by a lack of consensus regarding the selection of the most promising sites or genomic regions to target for analyses. The resulting heterogeneity presents challenges to researchers attempting to study the effects of early life stress on *OXTR* DNA methylation (8). We thus characterized the DNA methylation landscape of two putative functionally important regions, the entire CpG island in the gene promoter and a putative enhancer element in intron 3, to derive recommendations for future research. We report four major findings.

First, by using highly sensitive bisulfite sequencing (22), we point out 166 CpG sites with desirable statistical properties for research of interindividual differences (see assay sensitivity in Supplementary Table 1 for details). CpG sites with insufficient variance and low mean DNA methylation values (1%) can be disregarded in future studies.

Second, we used complexity reduction methods to derive clusters of co-methylated CpG sites. The majority of previous studies targeted only a few individual CpG sites either in the MT2 area (e.g. 18) or in the protein-coding region of exon 3 (e.g. 13), and recent efforts to find a consensus regarding the best sites to target for DNA methylation analysis remain challenging (8). Since the entire *OXTR* CpG island may be of potential functional relevance, the current study goes beyond the often-cited areas (i.e. MT2 and exon 3) and investigated all CpG sites of the CpG island. Instead of averaging across a certain number of CpG sites, a more informative data reduction strategy was used to detect patterns of interrelation using a data-driven method (38). Indeed, performing two different cluster approaches (agglomerative hierarchical clustering analysis and DBSCAN) led to the detection of an intrinsic structure to the *OXTR* gene. Two clusters, one spanning CpG sites within intron 1 and exon 2, and another spanning the protein-coding region of exon 3 were obtained, which contain largely redundant information. In addition, the DBSCAN procedure suggests no further clusters for the remaining CpG sites. Within the two clusters, intercorrelations range from *r*=.76 to *r*=.81, which seems to justify forming a mean DNA methylation value of clustered CpG sites, or to investigate only a few CpG sites to infer the mean DNA methylation value of one cluster. At other sites, CpG-wise inference should be performed.

Third, we report associations between DNA methylation, childhood trauma and gene expression, respectively. PLS implicates the same clusters (intron1/exon2 and protein-coding region of exon 3) together with several CpG sites within MT2 area (Figure 4) to contribute most strongly to the latent dimension predicting gene expression and CTQ scores. Thus, CpG sites in these segments seem to be statistically suitable for clinical between person studies. However, since there was low crossvalidated variance explained for the prediction of gene expression in the current study, the biological meaningfulness of these CpG sites remains unclear. Unfortunately, studies investigating consequences of altered DNA methylation on gene expression in human peripheral tissue are rare although the assessment of gene expression is necessary in order to gain a better insight into functional consequences of potential DNA methylation changes (39). A recent investigation in a translational animal model suggests that only DNA methylation in the MT2 area is predictive of *Oxtr* expression (40). However, investigations in human brain tissue revealed contradictory findings (41,42). As *OXTR* expression levels in peripheral tissue seem to be very low compared to brain tissue (https://www.proteinatlas.org/ENSG00000180914-OXTR), the functional relevance of *OXTR* DNA methylation remains be shown in human (post-mortem) brain tissue in the future.

Fourth, we observed extremely high mother-child correspondence between correlation patterns within the clusters described above (intron1/exon 2: *r*=.87 and exon3: *r*=.93), but only modest mother-child correspondence outside these clusters (r=.32). Moreover, mother-child correspondence in cluster solutions is very high as demonstrated by an overlap in CpG-to-cluster assignment of 98-100% depending on the clustering approach. This correspondence, which has hardly been studied so far, indicates a high generalizability of correlation patterns within these two clusters for different developmental stages (adult *vs*. child) and different tissues (PBMC *vs*. UMBC). In contrast, correlation patterns outside these regions exhibit low generalizability, potentially reflecting more dynamic DNA methylation-related processes which are relatively disjunct for different CpG sites.

It should be taken into account that while the present study has above-median sample size relative to prior studies on *OXTR* DNA methylation (8), its statistical power might be limited for some effect sizes which could still be considered theoretically meaningful. As this study was a secondary analysis of existing data, we did not perform power analyses for determination of the sample size but calculated the power post-hoc. Most descriptive effect sizes for associations of DNA methylation at single CpGs were small and we had limited power given our sample size of 110 mothers to detect effect sizes in this range (Supplementary Figure 13). Notably, the main focus of this paper was not to identify single significant CpGs or regions of CpG sites in association with CM or mRNA expression but rather to reduce the amount of CpG sites into functionally relevant clusters. Furthermore, results of the specific CpG sites were not consistent for different measures of CM (group status and CTQ). Thus, specific p-values or FDR values obtained from our analyses may only be informative in the light of our sample, but effect sizes, average differences in DNA methylation and p-values may be suggestive for highlighting specific regions of interest, together with results from cluster- and component-based analyses. Strikingly, effect sizes for associations between DNA methylation of single CpGs and CM were statistically small to medium (β_max_=|.48| for CTQ, Hedges’ g_max_=|0.67| for the group comparison). Given the small unstandardized group differences (maximum mean difference of 4% points) and confidence intervals being close to zero, the biological relevance of DNA methylation differences at single CpG sites remains unclear (see Supplementary Table 1 for details).

To conclude, the highly sensitive method for DNA methylation analysis, the application of various novel statistical methods to achieve complexity reduction as well as the parallel assessment of both DNA methylation and gene expression patterns reflect first steps towards development of best practices for the choice of CpG sites to study *OXTR* DNA methylation. The present study highlights clusters of CpG sites (intron1/exon2 and protein-coding region of exon3) that show desirable statistical properties for between person studies and are tentatively associated with childhood trauma and gene expression. Furthermore, we provide a Companion Web Application to guide future studies in their choice of CpG sites.

## Acknowledgements

The data has been collected within the BMBF-funded study “My childhood – Your childhood” (funding number: 01KR1304A). We are grateful for the support of the whole maternity ward staff at Ulm University Hospital, in particular Prof. Dr. med. Frank Reister. We also would like to thank the whole “My Childhood – Your Childhood” team. The DNA methylation analysis pipeline was developed under the scope of the BMBF-funded study “ProChild” (funding number: 01KR1805B).

## References

1. Feldman R, Monakhov M, Pratt M, Ebstein RP. Oxytocin Pathway Genes: Evolutionary Ancient System Impacting on Human Affiliation, Sociality, and Psychopathology. Biol Psychiatry. 2016;79:174–84.

2. Hammock EAD. Developmental Perspectives on Oxytocin and Vasopressin. Neuropsychopharmacology. 2015;40:24–42.

3. Heinrichs M, von Dawans B, Domes G. Oxytocin, vasopressin, and human social behavior. Front Neuroendocrinol. 2009;30:548–57.

4. Baribeau DA, Anagnostou E. Oxytocin and vasopressin: linking pituitary neuropeptides and their receptors to social neurocircuits. Front Neurosci. 2015;9:335.

5. Grinevich V, Stoop R. Interplay between Oxytocin and Sensory Systems in the Orchestration of Socio-Emotional Behaviors. Neuron. 2018;99:887–904.

6. Kompier NF, Keysers C, Gazzola V, Lucassen PJ, Krugers HJ. Early Life Adversity and Adult Social Behavior: Focus on Arginine Vasopressin and Oxytocin as Potential Mediators. Front Behav Neurosci. 2019;13:143.

7. Perkeybile AM, Carter CS, Wroblewski KL, Puglia MH, Kenkel WM, Lillard TS, et al. Early nurture epigenetically tunes the oxytocin receptor. Psychoneuroendocrinology. 2019;99:128–36.

8. Ellis BJ, Horn AJ, Carter CS, van IJzendoorn MH, Bakermans-Kranenburg MJ. Developmental programming of oxytocin through variation in early-life stress: Four meta-analyses and a theoretical reinterpretation. Clin Psychol Rev. 2021;86:101985.

9. Kraaijenvanger EJ, He Y, Spencer H, Smith AK, Bos PA, Boks MPM. Epigenetic variability in the human oxytocin receptor (OXTR) gene: A possible pathway from early life experiences to psychopathologies. Neurosci Biobehav Rev. 2019;96:127–42.

10. Kumsta R, Hummel E, Chen FS, Heinrichs M. Epigenetic regulation of the oxytocin receptor gene: implications for behavioral neuroscience. Front Neurosci. 2013;7:83.

11. Kusui C, Kimura T, Ogita K, Nakamura H, Matsumura Y, Koyama M, et al. DNA Methylation of the Human Oxytocin Receptor Gene Promoter Regulates Tissue-Specific Gene Suppression. Biochem Biophys Res Commun. 2001;289:681–6.

12. Needham BL, Smith JA, Zhao W, Wang X, Mukherjee B, Kardia SLR, et al. Life course socioeconomic status and DNA methylation in genes related to stress reactivity and inflammation: The multi-ethnic study of atherosclerosis. Epigenetics. 2015;10:958–69.

13. Unternaehrer E, Meyer AH, Burkhardt SCA, Dempster E, Staehli S, Theill N, et al. Childhood maternal care is associated with DNA methylation of the genes for brain-derived neurotrophic factor (*BDNF*) and oxytocin receptor (*OXTR*) in peripheral blood cells in adult men and women. Stress. 2015;18:451–61.

14. Fujisawa TX, Nishitani S, Takiguchi S, Shimada K, Smith AK, Tomoda A. Oxytocin receptor DNA methylation and alterations of brain volumes in maltreated children. Neuropsychopharmacology. 2019;44:2045–53.

15. Gouin JP, Zhou QQ, Booij L, Boivin M, Côté SM, Hébert M, et al. Associations among oxytocin receptor gene (OXTR) DNA methylation in adulthood, exposure to early life adversity, and childhood trajectories of anxiousness. Sci Rep. 2017;7:7446.

16. Smearman EL, Almli LM, Conneely KN, Brody GH, Sales JM, Bradley B, et al. Oxytocin Receptor Genetic and Epigenetic Variations: Association With Child Abuse and Adult Psychiatric Symptoms. Child Dev. 2016;87:122–34.

17. Robakis TK, Zhang S, Rasgon NL, Li T, Wang T, Roth MC, et al. Epigenetic signatures of attachment insecurity and childhood adversity provide evidence for role transition in the pathogenesis of perinatal depression. Transl Psychiatry. 2020;10:48.

18. Krol KM, Moulder RG, Lillard TS, Grossmann T, Connelly JJ. Epigenetic dynamics in infancy and the impact of maternal engagement. Sci Adv. 2019;5:eaay0680.

19. King L, Robins S, Chen G, Yerko V, Zhou Y, Nagy C, et al. Perinatal depression and DNA methylation of oxytocin-related genes: a study of mothers and their children. Horm Behav. 2017;96:84–94.

20. Ramo-Fernández L, Gumpp AM, Boeck C, Krause S, Bach AM, Waller C, et al. Associations between childhood maltreatment and DNA methylation of the oxytocin receptor gene in immune cells of mother–newborn dyads. Transl Psychiatry. 2021;11:449.

21. Inoue T, Kimura T, Azuma C, Inazawa J, Takemura M, Kikuchi T, et al. Structural organization of the human oxytocin receptor gene. J Biol Chem. 1994;269:32451–6.

22. Moser DA, Müller S, Hummel EM, Limberg AS, Dieckmann L, Frach L, et al. Targeted bisulfite sequencing: A novel tool for the assessment of DNA methylation with high sensitivity and increased coverage. Psychoneuroendocrinology. 2020;120:104784.

23. Bader K, Hänny C, Schäfer V, Neuckel A, Kuhl C. Childhood Trauma Questionnaire – Psychometrische Eigenschaften einer deutschsprachigen Version. Z Für Klin Psychol Psychother. 2009;38:223–30.

24. Berstein DP, Fink L. Childhood trauma questionnaire: a retrospective self-report manual. San Antonio, TX: Psychological Corporation, 1998.

25. Leitão E, Beygo J, Zeschnigk M, Klein-Hitpass L, Bargull M, Rahmann S, et al. (2018). Locus-Specific DNA Methylation Analysis by Targeted Deep Bisulfite Sequencing. In: Jeltsch A, Rots M (eds). Epigenome Editing: Methods in Molecular Biology. Humana Press: New York, 1767, pp 351–366.

26. Rahmann S, Beygo J, Kanber D, Martin M, Horsthemke B, Buiting K. Amplikyzer: Automated methylation analysis of amplicons from bisulfite flowgram sequencing. PeerJ Preprints. 2013;1:e122v2.

27. Andersen CL, Jensen JL, Ørntoft TF. Normalization of Real-Time Quantitative Reverse Transcription-PCR Data: A Model-Based Variance Estimation Approach to Identify Genes Suited for Normalization, Applied to Bladder and Colon Cancer Data Sets. Cancer Res. 2004;64:5245–50.

28. Hahsler M, Piekenbrock M, Doran D. dbscan: Fast Density-Based Clustering with R. J Stat Softw. 2019;91:1–30.

29. Kuhn M, Wing J, Weston S, Williams A, Keefer C, Engelhardt A, et al. (2021). caret: Classification and Regression Training. R package version v6.0-90. https://CRAN.R-project.org/package=caret.

30. Liland KH, Mevik B-H, Wehrens R, Hiemstra P. (2021). pls: Partial Least Squares and Principal Component Regression. R package version v2.8-0. https://CRAN.R-project.org/package=pls.

31. Chén OY, Crainiceanu C, Ogburn EL, Caffo BS, Wager TD, Lindquist MA. High-dimensional multivariate mediation with application to neuroimaging data. Biostatistics. 2018;19:121–36.

32. Geuter S, Reynolds Losin EA, Roy M, Atlas LY, Schmidt L, Krishnan A, et al. Multiple Brain Networks Mediating Stimulus–Pain Relationships in Humans. Cereb Cortex. 2020;30:4204–19.

33. Ritchie ME, Phipson B, Wu D, Hu Y, Law CW, Shi W, et al. limma powers differential expression analyses for RNA-sequencing and microarray studies. Nucleic Acids Res. 2015;43:e47.

34. Peters TJ, Buckley MJ, Statham AL, Pidsley R, Samaras K, V Lord R, et al. De novo identification of differentially methylated regions in the human genome. Epigenetics Chromatin. 2015;8:6.

35. Chen Y, Pal B, Visvader JE, Smyth GK. Differential methylation analysis of reduced representation bisulfite sequencing experiments using edgeR. F1000Res. 2017;6:2055.

36. Hansen KD, Langmead B, Irizarry RA. BSmooth: from whole genome bisulfite sequencing reads to differentially methylated regions. Genome Biol. 2012;13:R83.

37. Lecompte V, Robins S, King L, Solomonova E, Khan N, Moss E, et al. Examining the role of mother-child interactions and DNA methylation of the oxytocin receptor gene in understanding child controlling attachment behaviors. Attach Hum Dev. 2021;23:37–55.

38. Lancaster K, Morris JP, Connelly JJ. Neuroimaging Epigenetics: Challenges and Recommendations for Best Practices. Neuroscience. 2018;370:88–100.

39. Jones MJ, Moore SR, Kobor MS. Principles and Challenges of Applying Epigenetic Epidemiology to Psychology. Annu Rev Pychol. 2018;69:459–85.

40. Danoff JS, Wroblewski KL, Graves AJ, Quinn GC, Perkeybile AM, Kenkel WM, et al. Genetic, epigenetic, and environmental factors controlling oxytocin receptor gene expression. Clin Epigenetics. 2021;13:23.

41. Almeida D, Fiori LM, Chen GG, Aouabed Z, Lutz P-E, Zhang T-Y, et al. Oxytocin receptor expression and epigenetic regulation in the anterior cingulate cortex of individuals with a history of severe childhood abuse. Psychoneuroendocrinology. 2022;136:105600.

