## Supplementary Figures for "The DNA methylation landscape of the human oxytocin receptor gene (*OXTR*): Recommendations for future research"

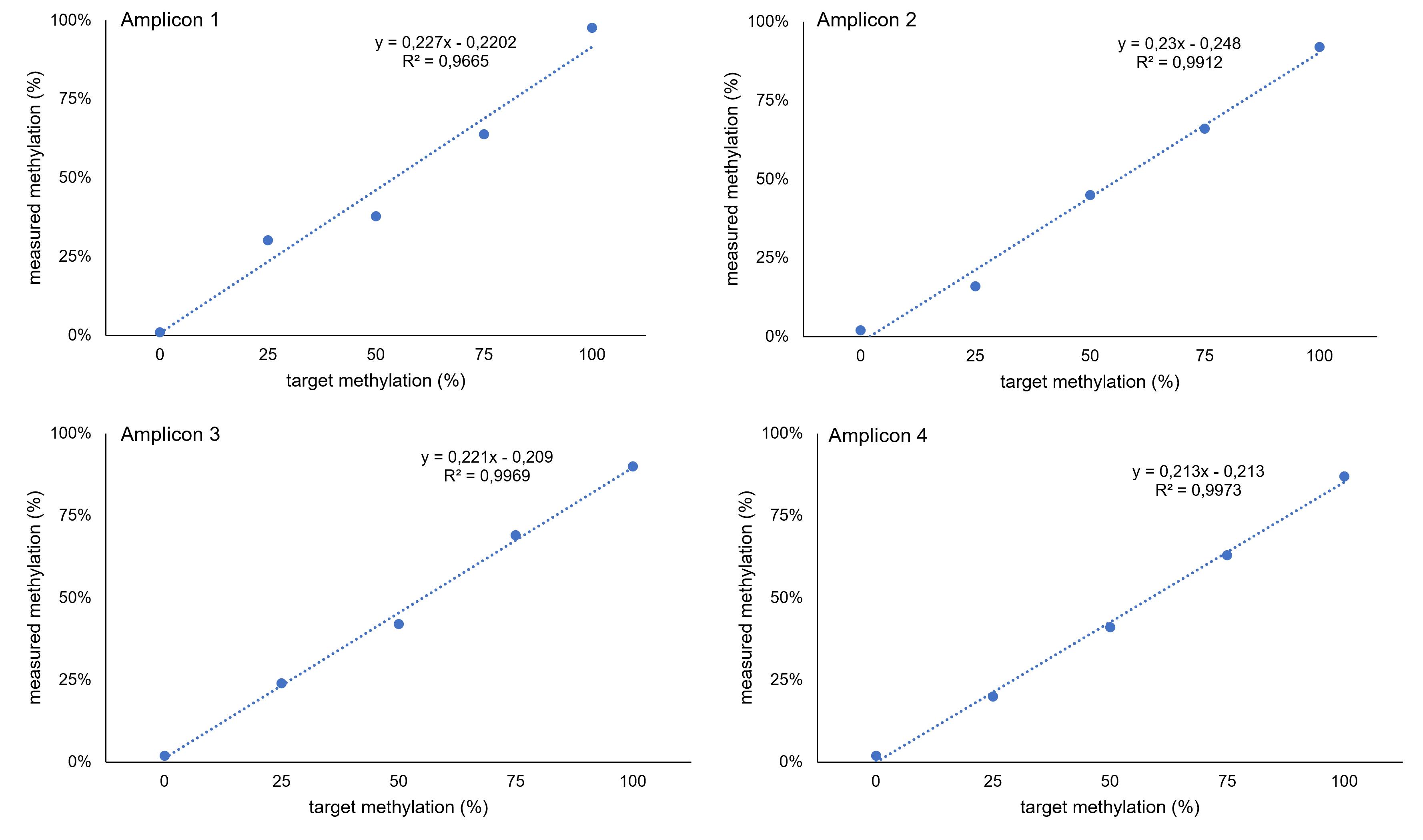

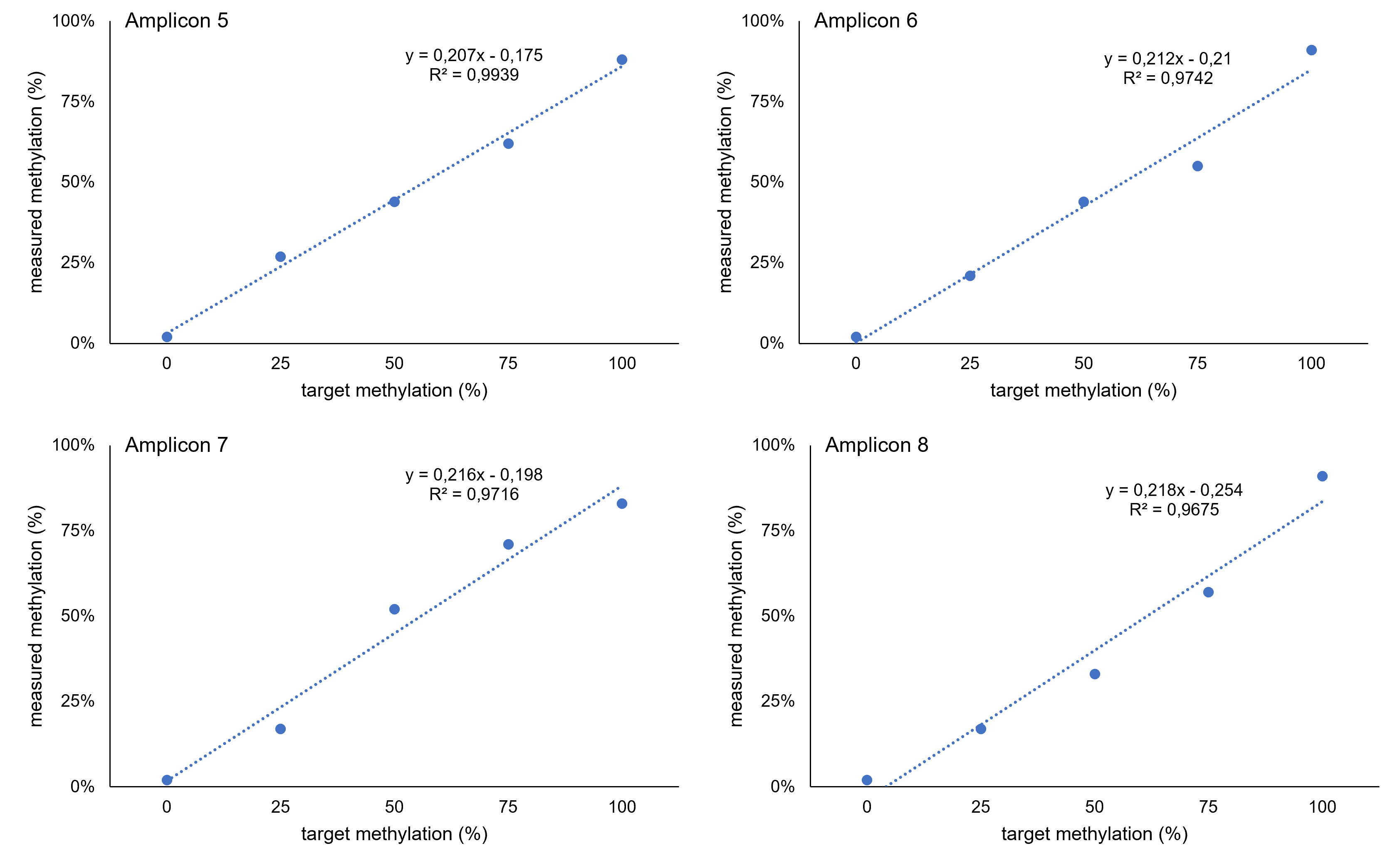

**Supplementary Figure 1: Standard curves for amplicons spanning the *OXTR* promoter region.** Linearity of the assays was tested by plotting expected to experimental methylation values of a serial diluted methylation standard. Several methylation standards with a known degree of methylation (calibration mixtures of 0%, 25%, 50%, 75% and 100% methylated DNA) for the first experiments on the MiSeq system were assayed. Results indicated no preferential DNA amplification in dependence of DNA methylation and results following a linear curve for all standards tested.

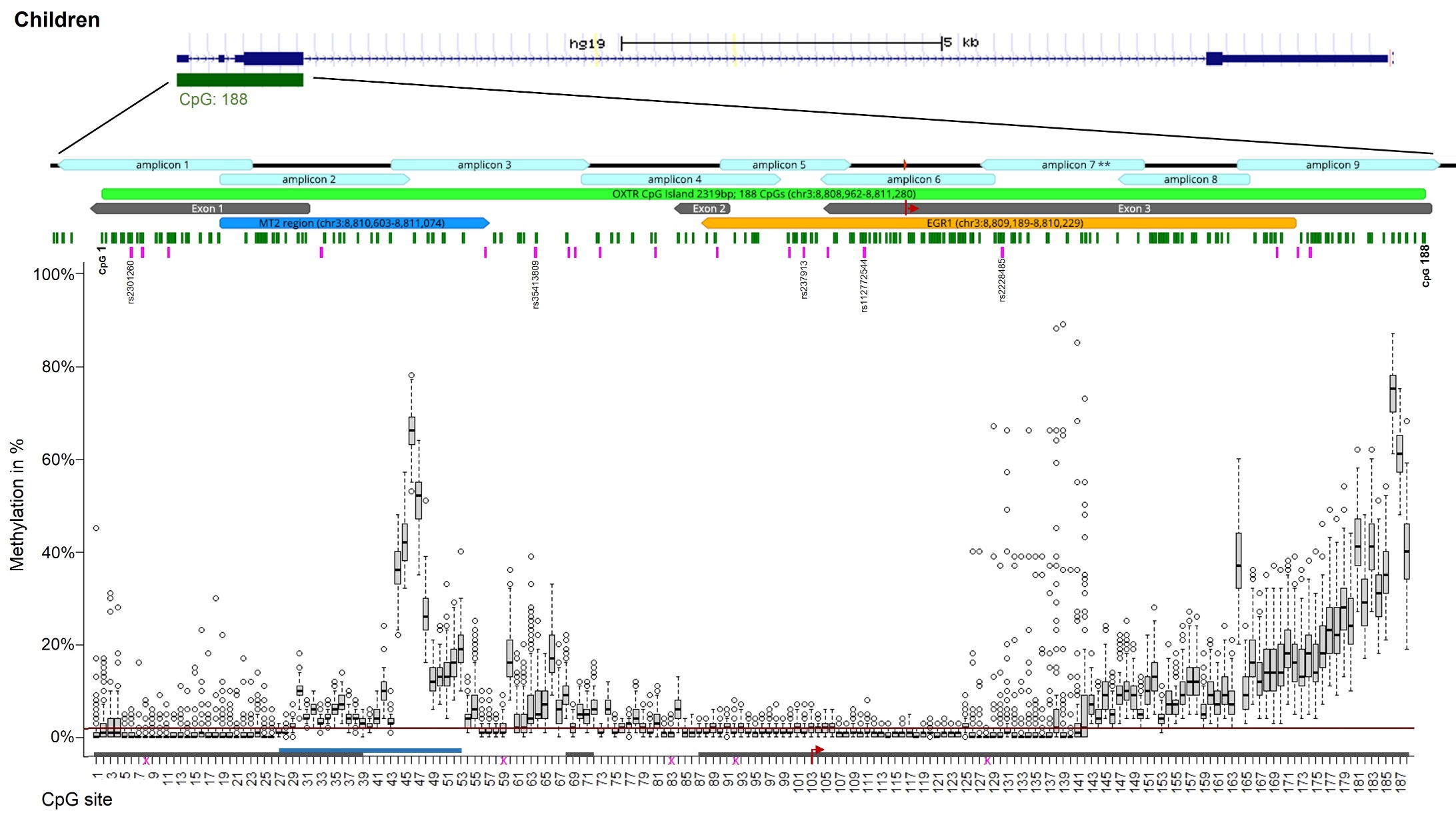

**Supplementary Figure 2. Chromosomal position of *OXTR* promoter region with corresponding DNA methylation levels (children).**
Chromosomal position of all fragments of the *OXTR* CpG island for children are illustrated using graphical outputs generated by the UCSC Genome Browser (https://genome.ucsc.edu) and Geneious Prime 2020 software (https://www.geneious.com). The gene is presented in 5’ -> 3’ orientation from left to right. Amplicons are highlighted with respect to their genomic orientation. All CpG sites are illustrated with green bars, CpG island in light green (with CpG1 and CpG 188 labelled), MT2 area in blue, and exons in grey colour (transcription start site is marked by the red arrow). A region including several transcription factor binding sites for EGRI (OREG1492306 according to the UCSC Genome Browser; also known as NGFIA) is indicated in yellow. Common SNPs with a minor allele frequency >5% according to the UCSC Genome Browser are illustrated with pink bars. Genomic regions of *OXTR* SNPs that introduce, disrupt or shift methylation sites are highlighted (these CpG sites were excluded from statistical analyses). Boxplots showing DNA methylation across the investigated CpG sites. The box covers the methylation data of each CpG site between the 25th to 75th quantile, the whiskers show the range of values falling within 1.5-fold the interquartile range. The horizontal line (red) represents the methylation detection limit for targeted deep bisulfite sequencing (*OXTR* promoter: 1,96%).
Note: ** The values in the range of amplicon 7 are not different due to epigenetic heterogeneity but rather due to technical variability.

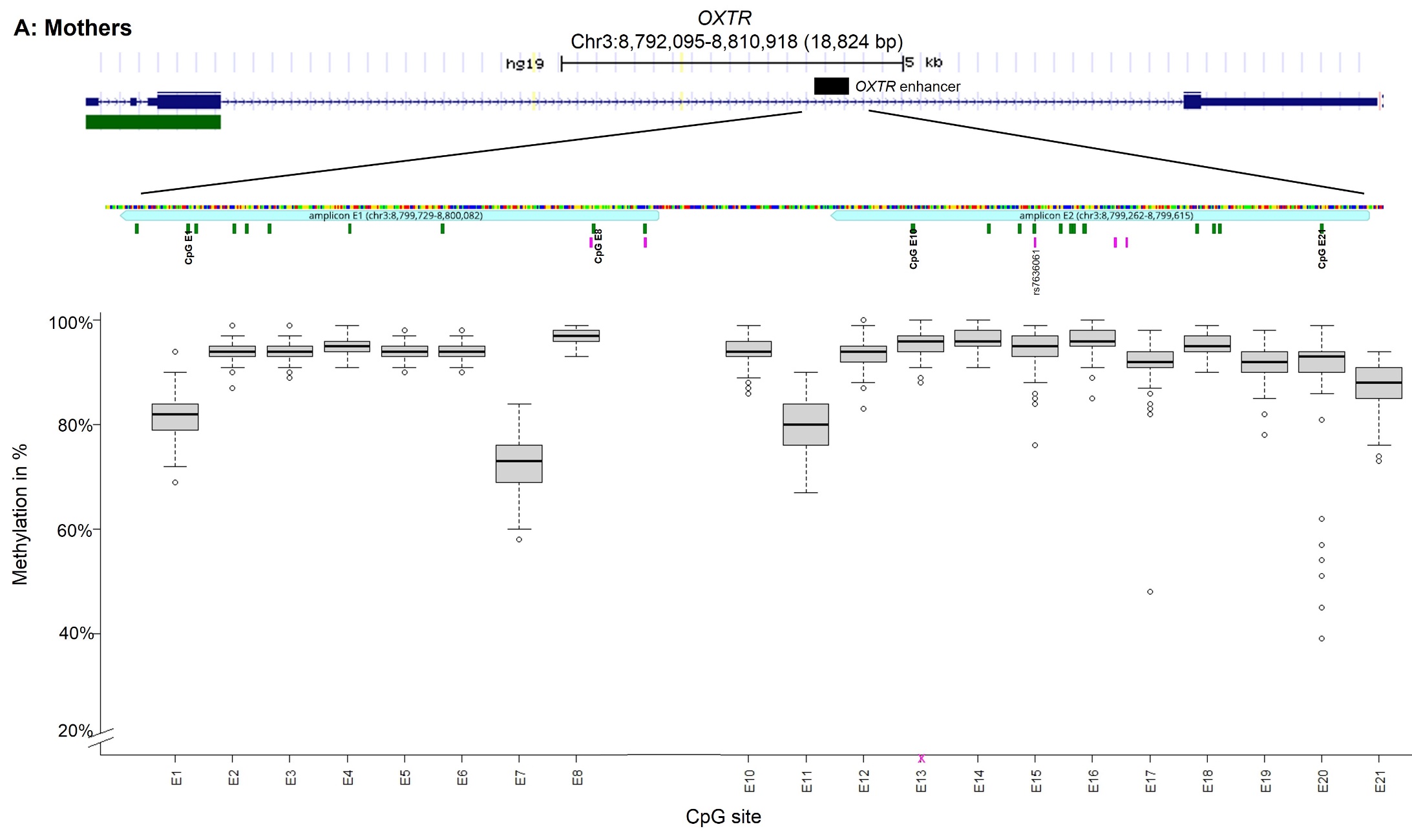

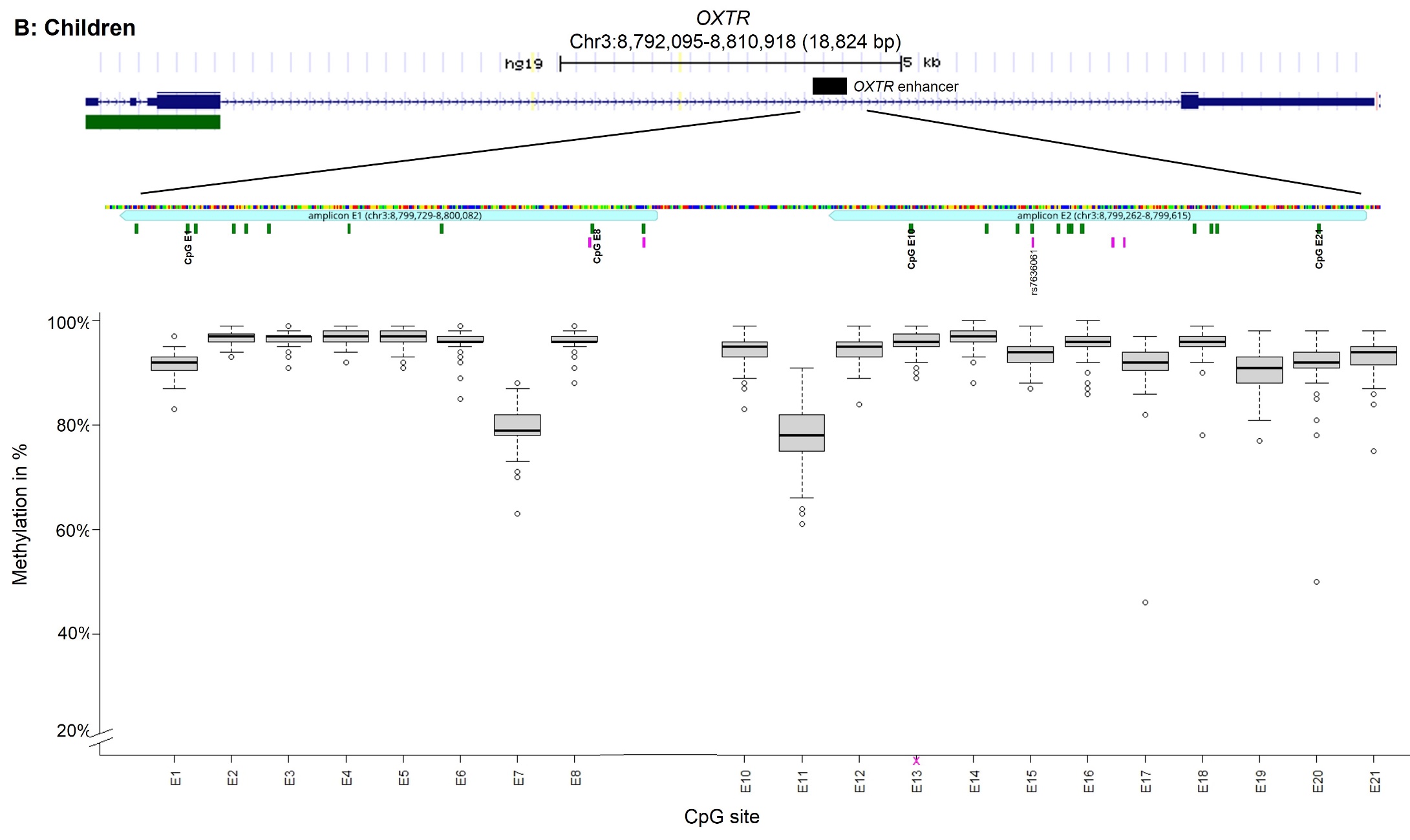

**Supplementary Figure 3: Chromosomal position of *OXTR* enhancer region with corresponding DNA methylation levels (A: mothers; B: children).** Chromosomal position of all fragments of a potential *OXTR* enhancer element, shown separately for mothers (A) and children (B) respectively, are illustrated using graphical outputs generated by the UCSC Genome Browser (https://genome.ucsc.edu) and Geneious Prime 2020 software (https://www.geneious.com). The gene is presented in 5’ -> 3’ orientation from left to right. Amplicons are highlighted with respect to their genomic orientation. All CpG sites are illustrated with green bars (with CpG E1 and CpG E21 labelled). Common SNPs with a minor allele frequency >5% according to the UCSC Genome Browser are illustrated with pink bars. Genomic region of *OXTR* SNPs that introduce, disrupt or shift methylation site is highlighted (this CpG site was excluded from statistical analyses). Boxplots showing DNA methylation across the investigated CpG sites. The box covers the methylation data of each CpG site between the 25th to 75th quantile, the whiskers show the range of values falling within 1.5-fold the interquartile range.

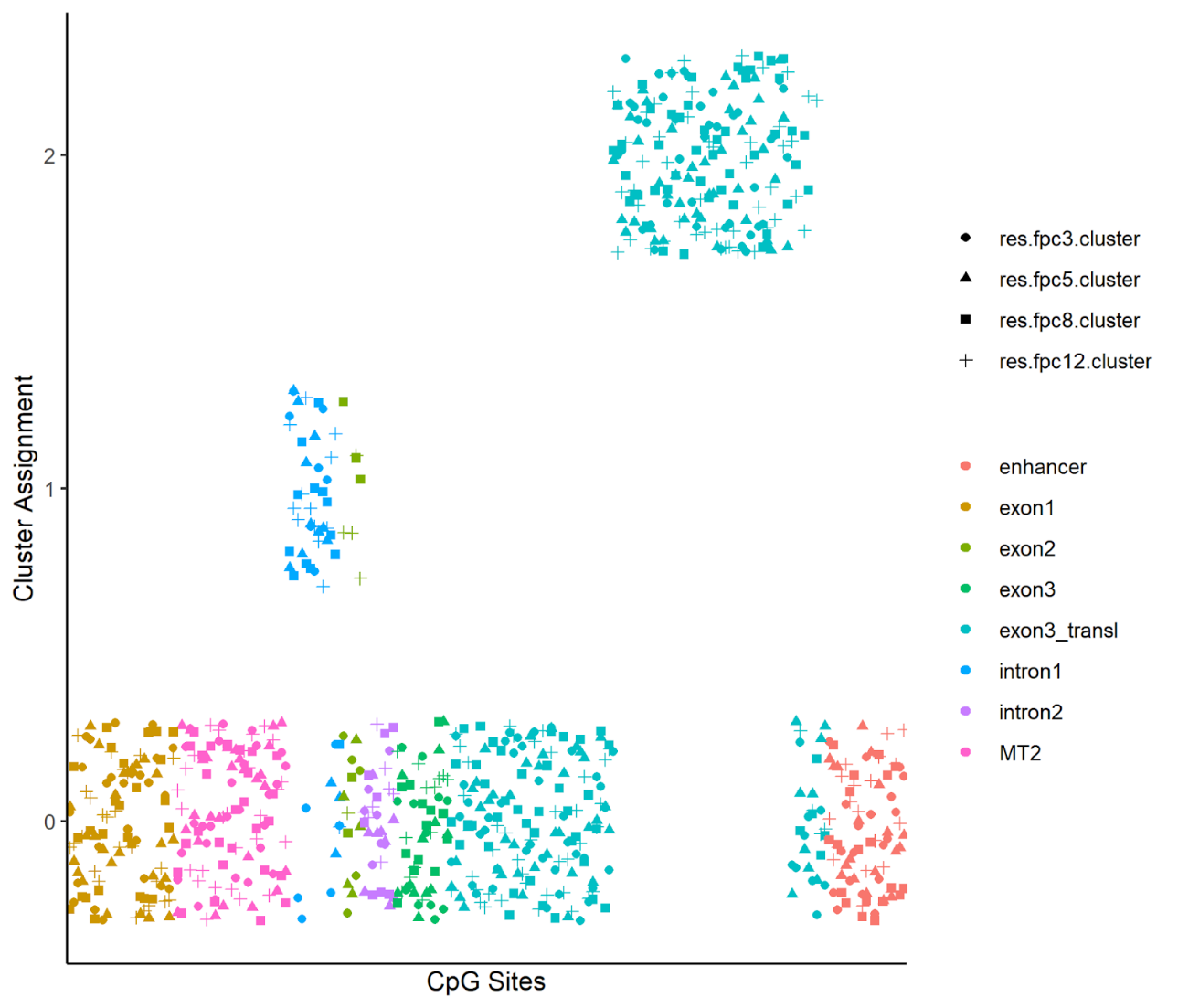

**Supplementary Figure 4. Clustering Solution of DBSCAN procedure in mothers.** Shapes distinguish between different hyperparameters. Possible values for cluster assignment were 0, 1, and 2. Datapoints were jittered around their cluster assignment value for visibility reasons. The cluster with assigned values zero represents the outlier category. Hyperparameter values:
res.fpc3.cluster: MinPts = 3, eps = 5; res.fpc5.cluster: MinPts = 5, eps = 5.8; res.fpc8.clsuter: MinPts = 8, eps = 7.7; res.fpc12: MinPts = 12, eps = 8.

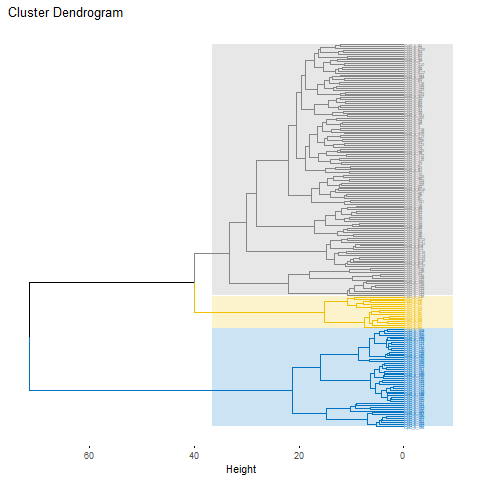

**Supplementary Figure 5. Hierarchical agglomerative clustering solution in children.** All CpG sites of the *OXTR* CpG island (comprising three exons and two introns) and a potential enhancer element were included.

 
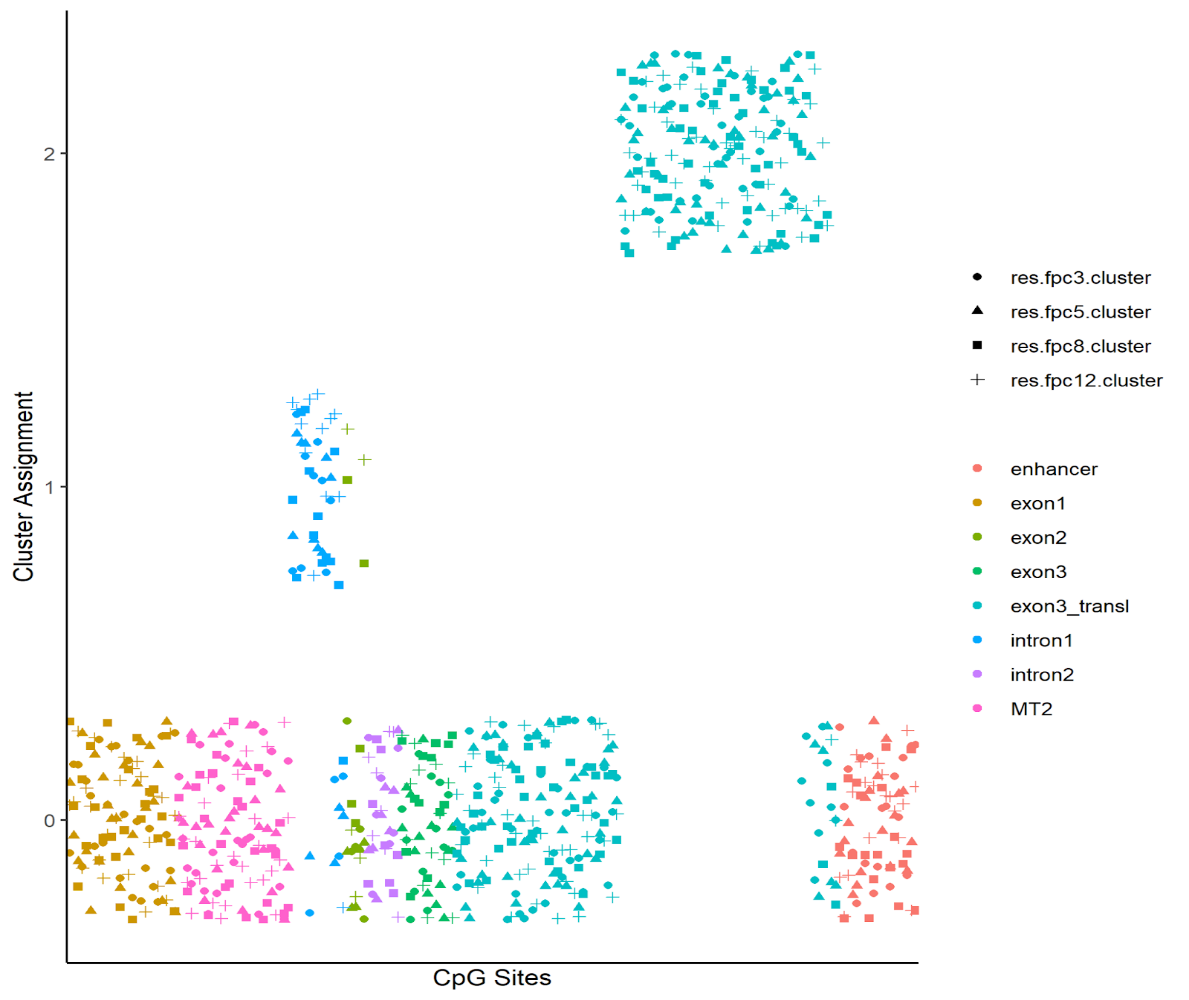

**Supplementary Figure 6. Clustering Solution of DBSCAN procedure in children.** Shapes distinguish between different hyperparameters. Possible values for cluster assignment were 0, 1, and 2. Datapoints were jittered around their cluster assignment value for visibility reasons. The cluster with assigned values zero represents the outlier category. Hyperparameter values: res.fpc3.cluster: MinPts = 3, eps = 5; res.fpc5.cluster: MinPts = 5, eps = 5.8; res.fpc8.clsuter: MinPts = 8, eps = 7.7; res.fpc12: MinPts = 12, eps = 8.

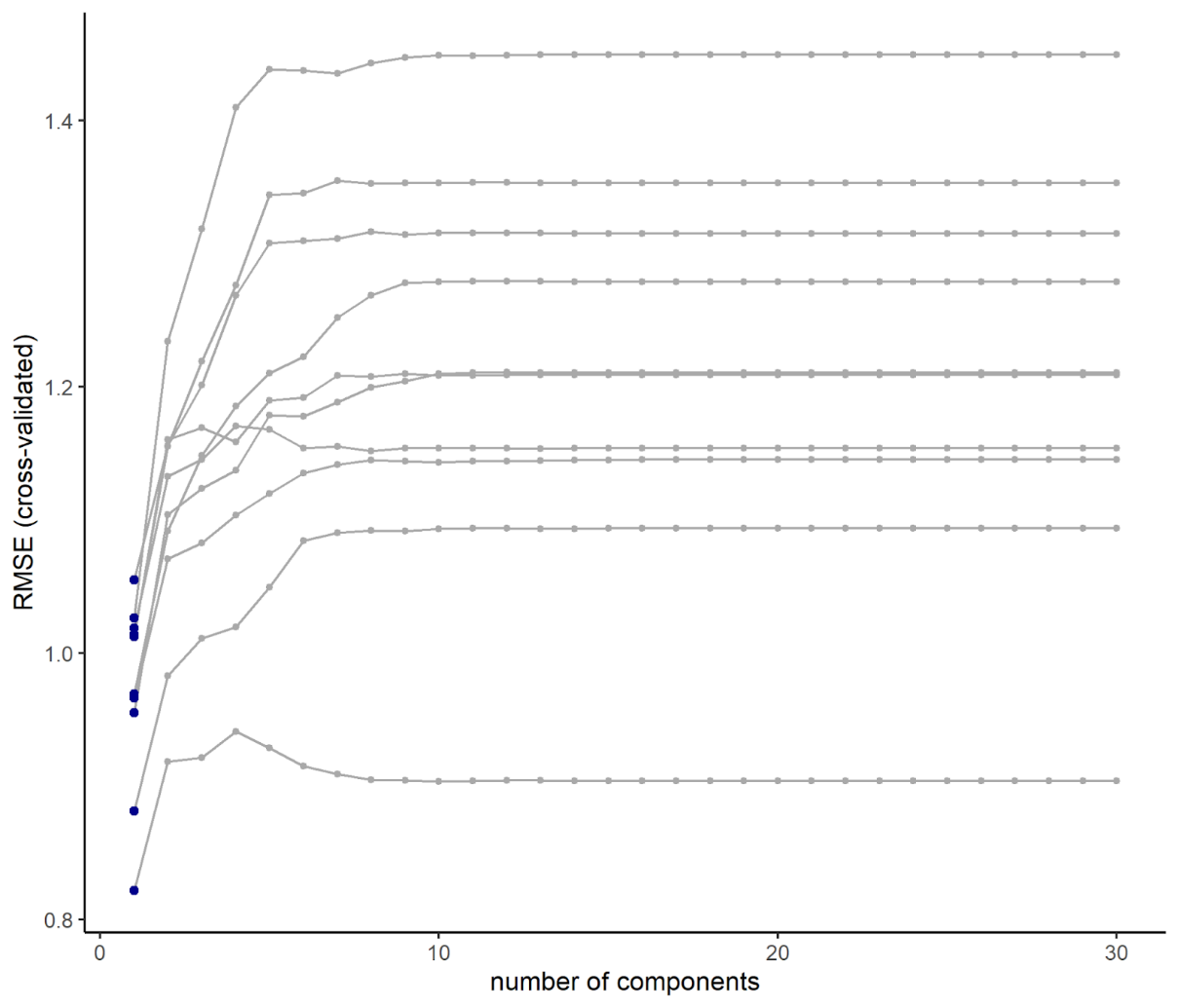

**Supplementary Figure 7. Hyperparameter tuning for gene expression.** Cross-validated root-mean-squared error (RMSE) as a function of number of components for the 2x5 = 10 outer cross-validation folds. The minimum RMSE for each fold is highlighted in blue.

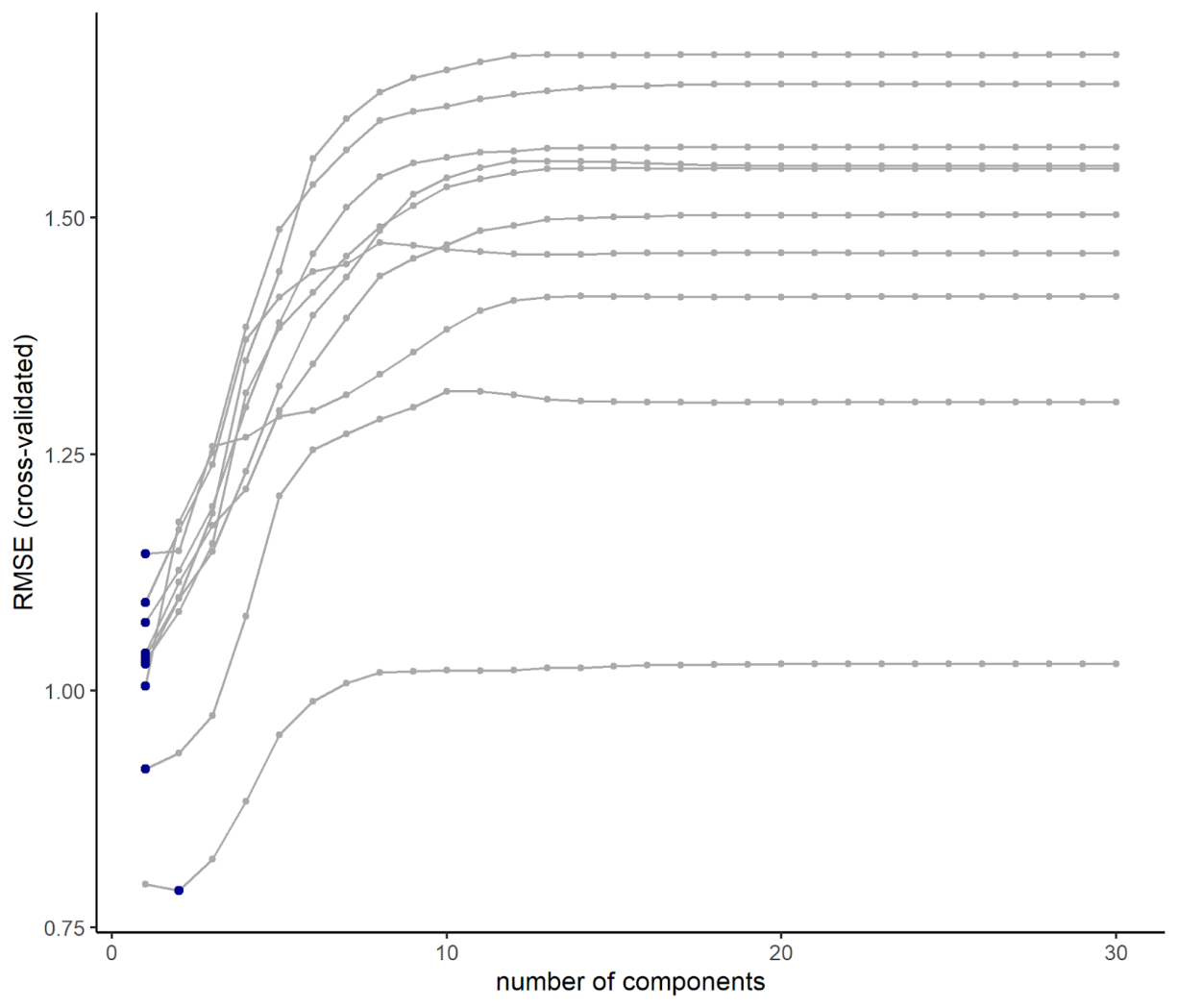

**Supplementary Figure 8. Hyperparameter tuning for childhood trauma questionnaire.** Cross-validated root-mean-squared error (RMSE) as a function of number of components for the 2x5 = 10 outer cross-validation folds. The minimum RMSE for each fold is highlighted in blue.

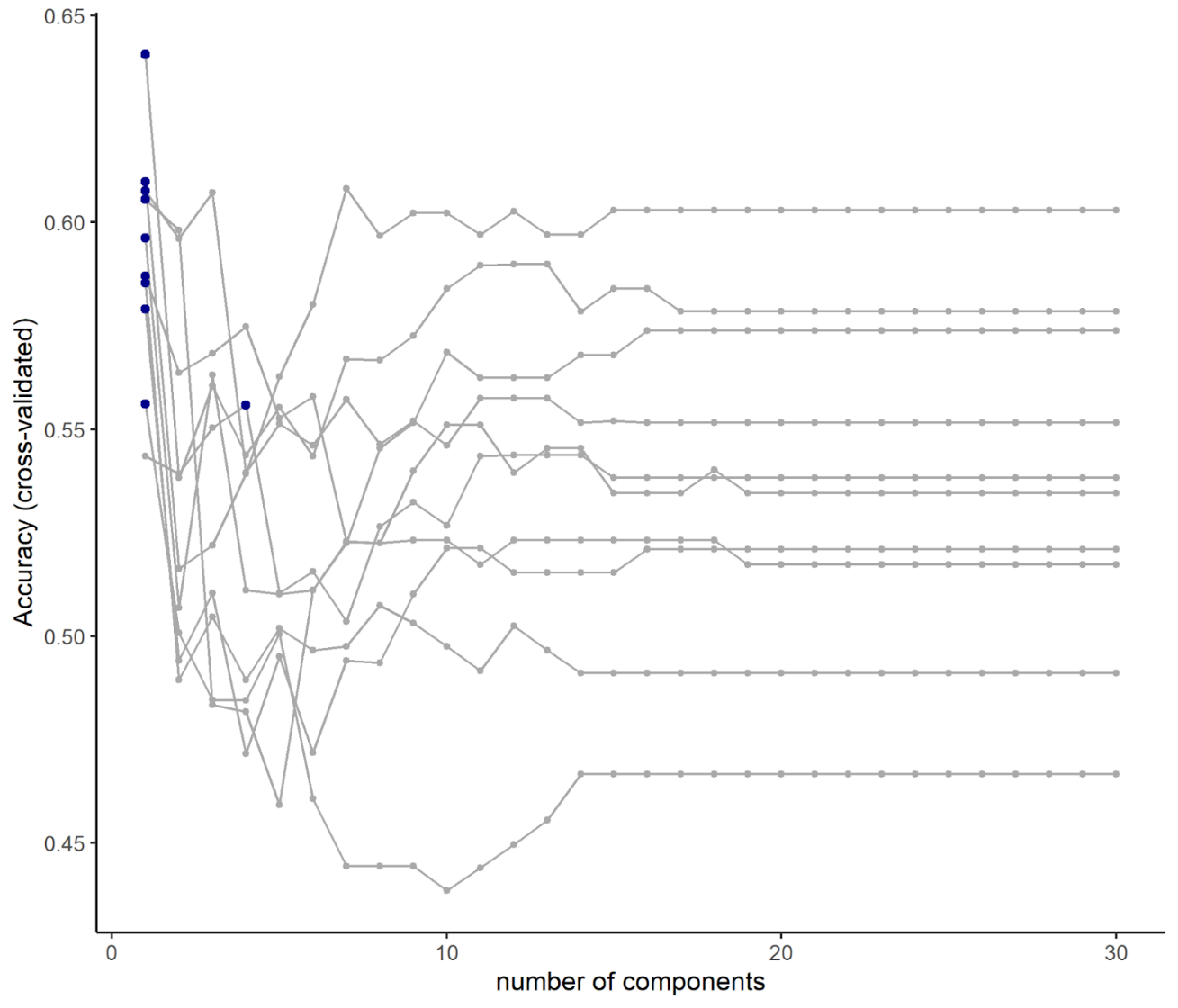

**Supplementary Figure 9. Hyperparameter tuning for binary trauma group discrimination.** Cross-validated classification accuracy as a function of number of components for the 2x5 = 10 outer cross-validation folds. The maximum accuracy for each fold is highlighted in blue.

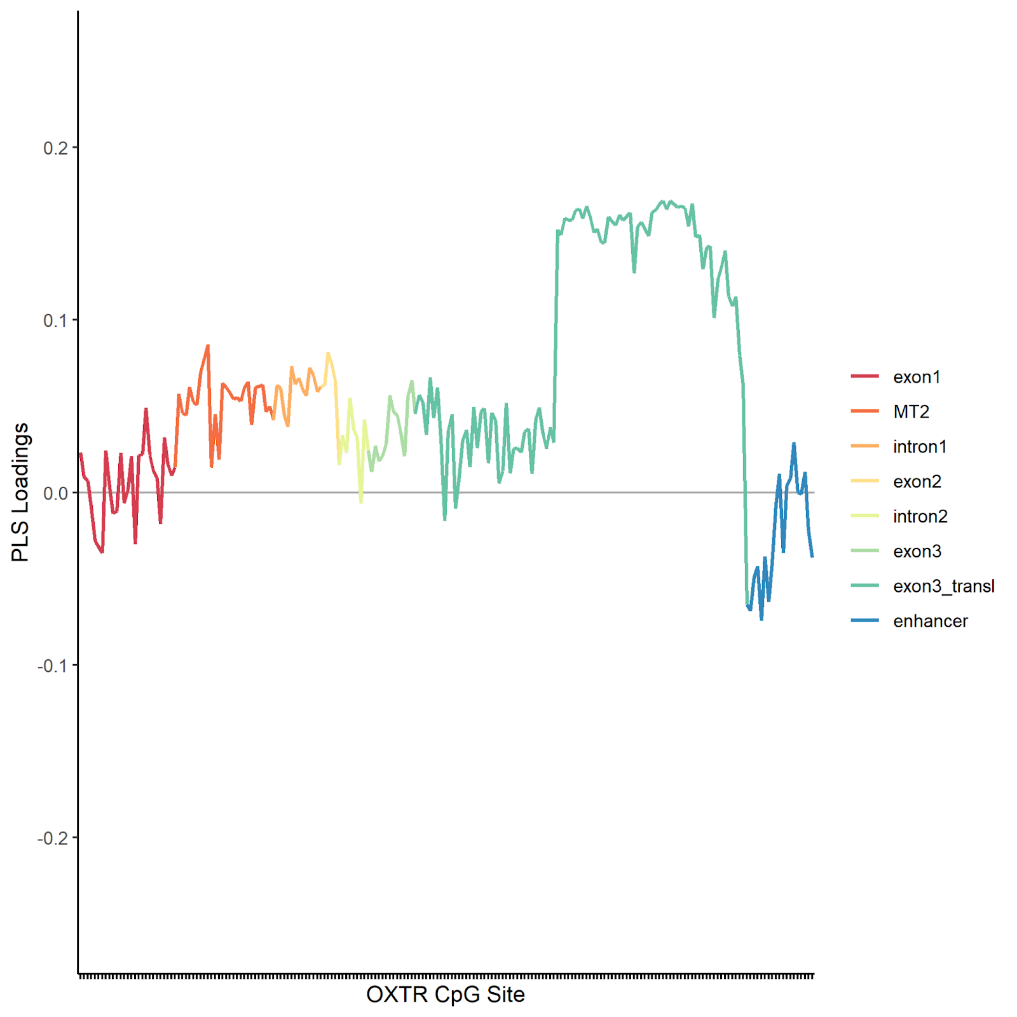

**Supplementary Figure 10. Partial Least Squares loadings of *OXTR* DNA methylation at single CpG sites for the prediction of binary trauma group.** Analyses included 166 CpG sites of the *OXTR* CpG island as predictors, after excluding thirty-six CpG sites due to insufficient variance.

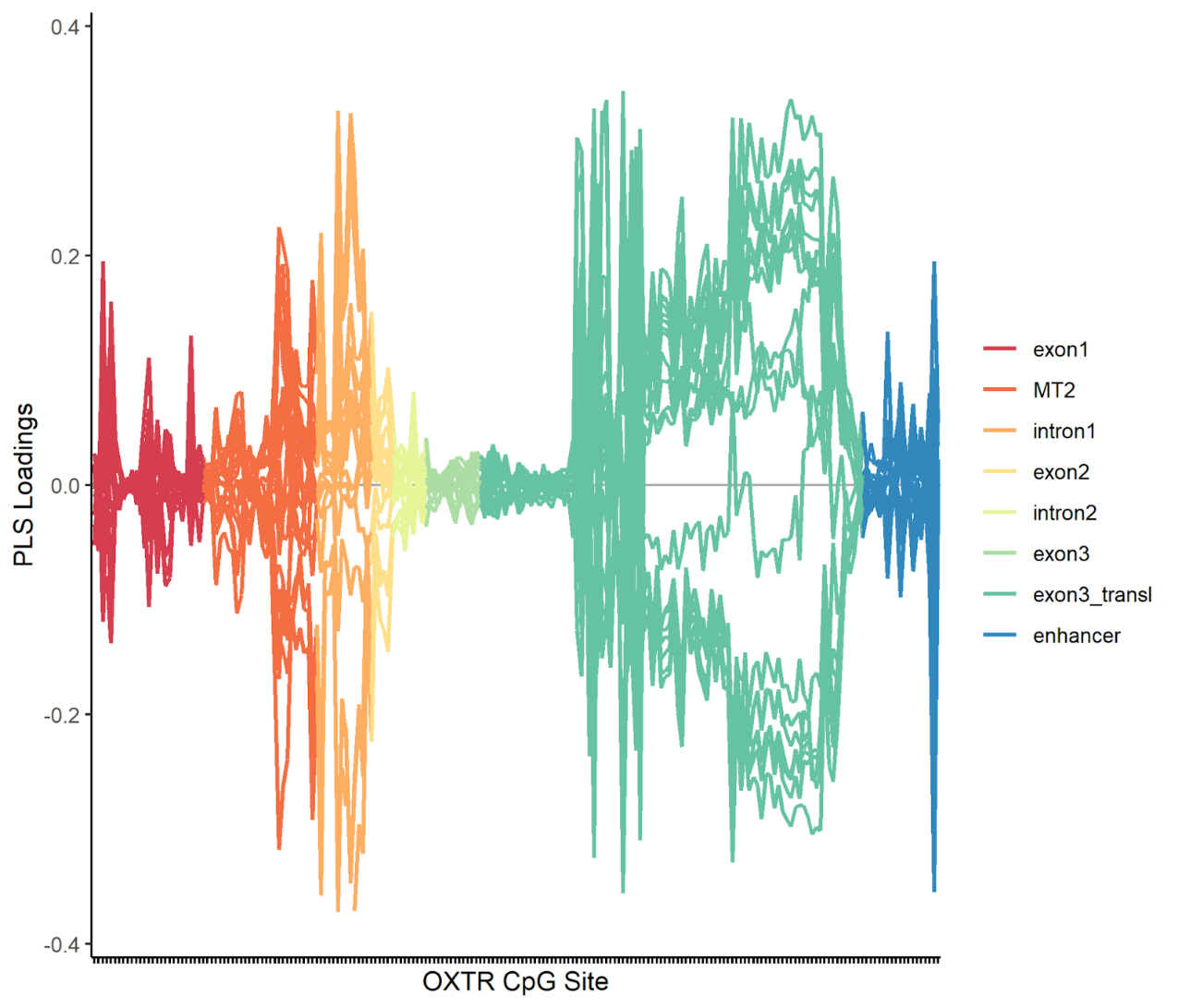

**Supplementary Figure 11. Partial Least Squares loadings of *OXTR* DNA methylation at single CpG sites for the prediction of randomly created variables (20 iterations).**

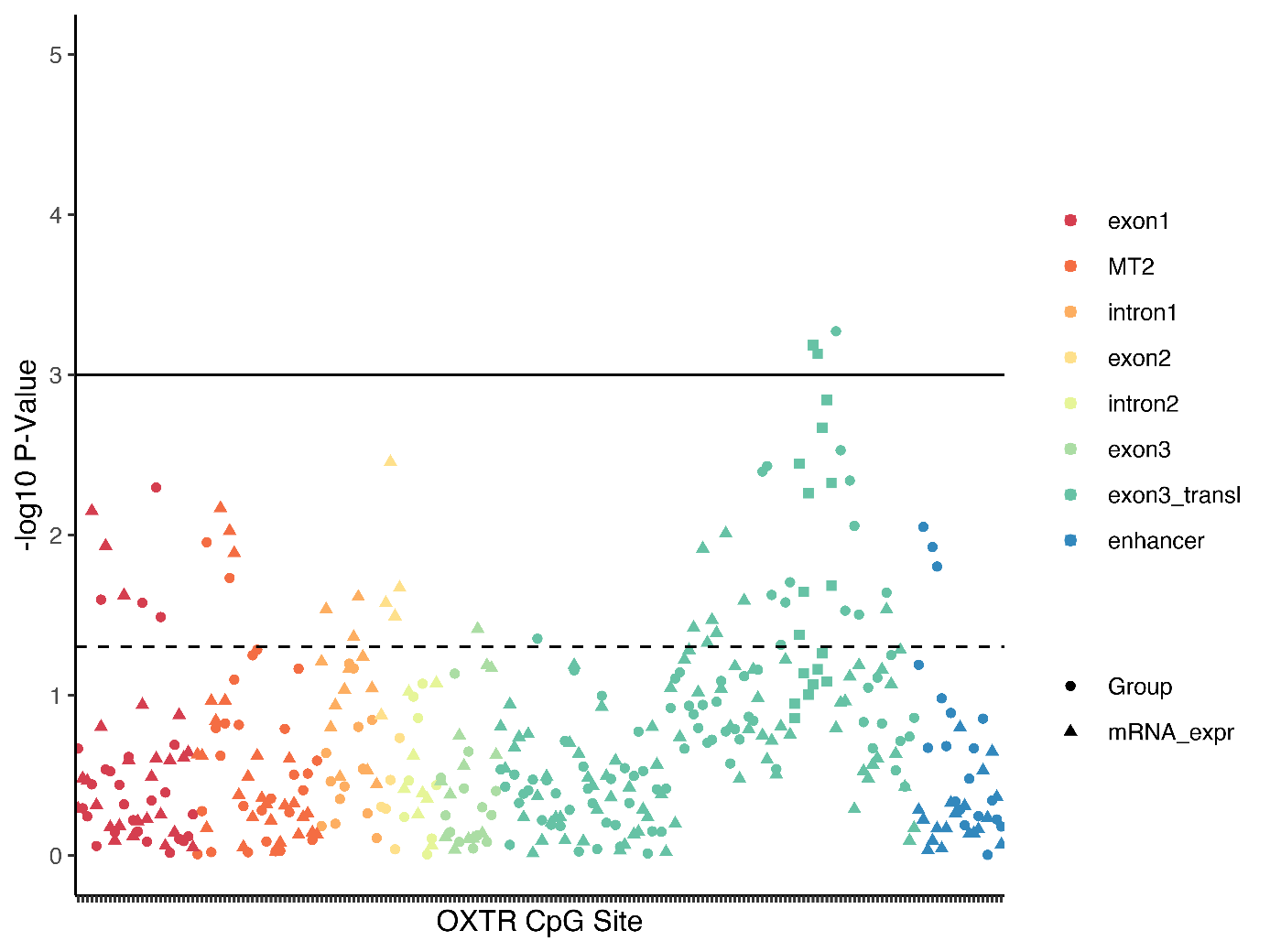

**Supplementary Figure 12. Associations of childhood maltreatment (group status) and *OXTR* mRNA expression with DNA methylation at single CpG sites.** Triangles depict the -log10 p-value of the association between DNA methylation and mRNA expression, circles depict the association with childhood maltreatment (CM) and squares indicate the DMR in association with CM. The dotted line indicates a nominal p-value of .05, whereas the solid line indicates an FDR of .05.

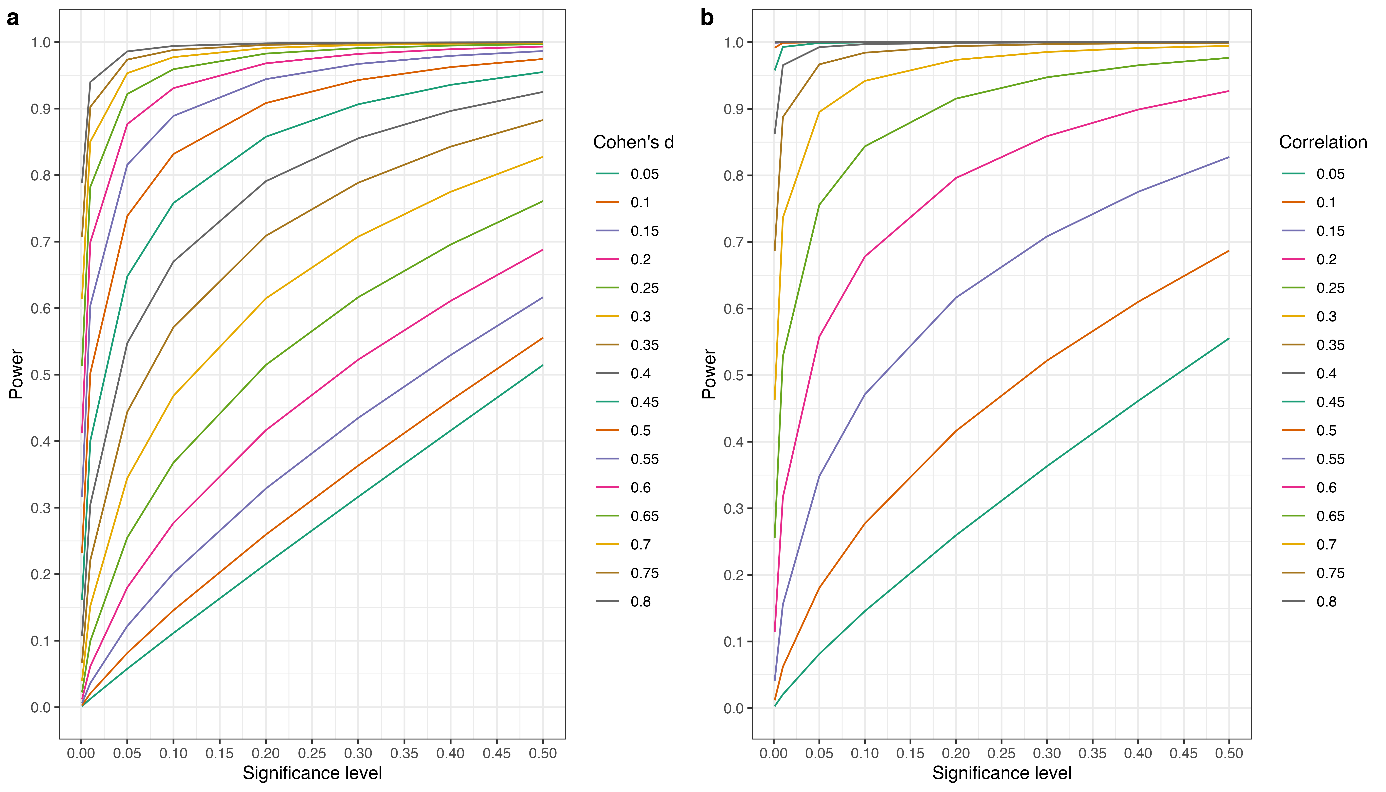

**Supplementary Figure 13. Power curves depending on varying effect sizes and alpha levels, but with fixed sample size. a** Analyses for two-sided independent t-tests with N=55 per group. **b** Analyses for correlation test (equivalent to standardised regression coefficients) with N=110.

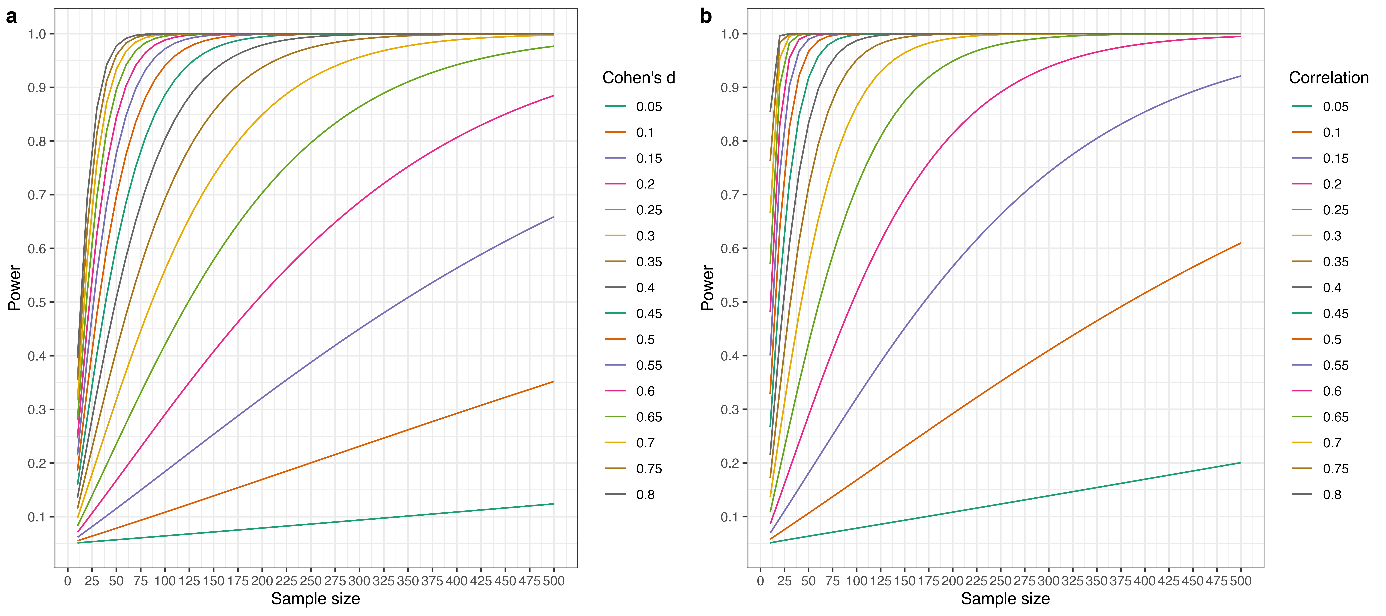

**Supplementary Figure 14. Power curves depending on varying effect sizes and sample sizes, but with fixed alpha level. a** Analyses for two-sided independent t-tests with α=.05. Sample size refers to N per group. **b** Analyses for correlation test (equivalent to standardised regression coefficients) with α=.05.

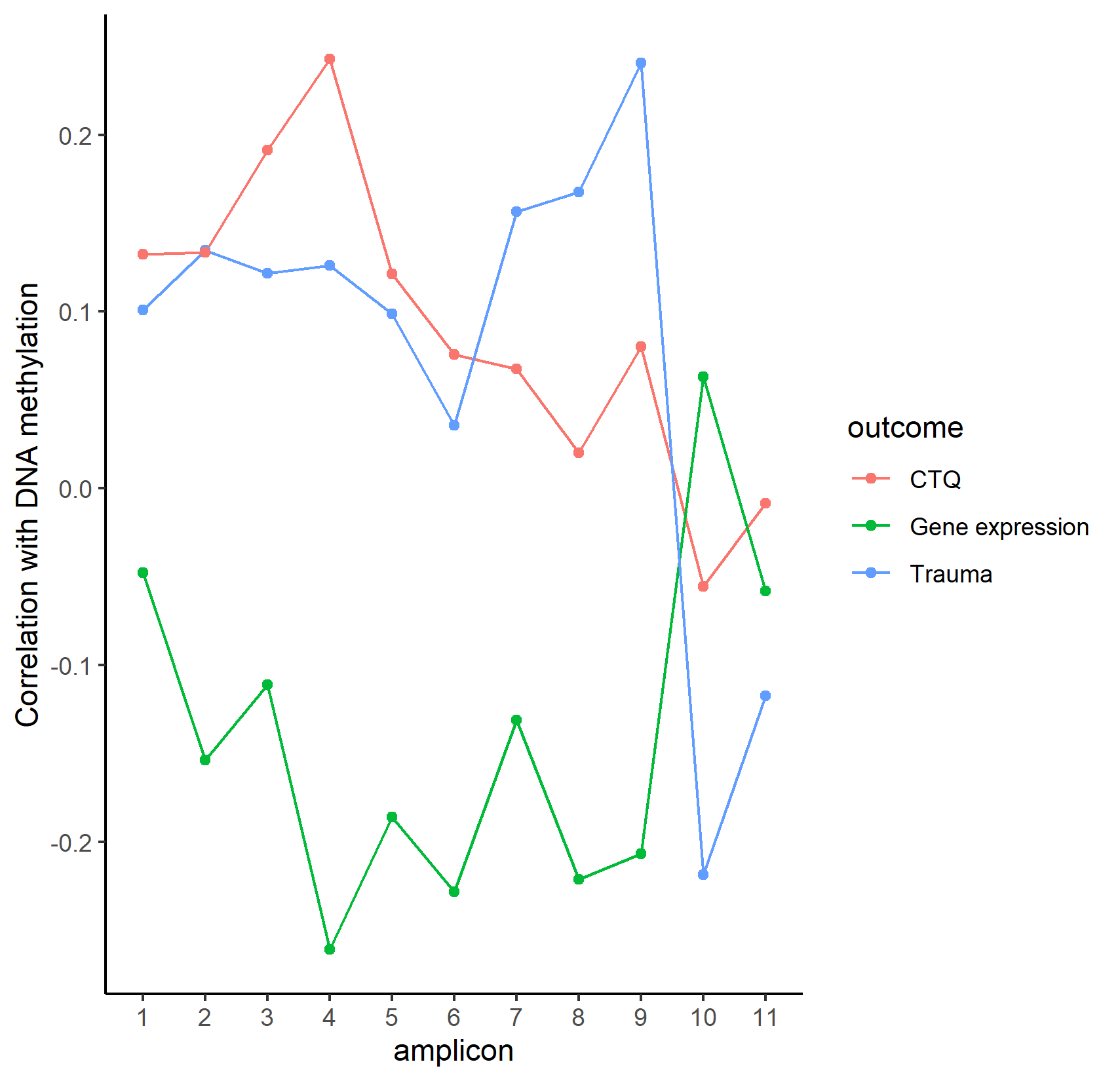

**Supplementary Figure 15. Correlation between average *OXTR* DNA methylation in the respective amplicons and the three external outcomes of the study.** CTQ = childhood trauma questionnaire.
